# Integrative Analysis of Spatial Transcriptome and Connectome by SpaCon

**DOI:** 10.64898/2025.12.11.693415

**Authors:** Haichao Qu, Wenyuan Li, Yao Fei, Shijie Zhao, Junwei Han, Cirong Liu

## Abstract

The brain’s structure and function arise from its complex molecular composition and neural connectivity, yet the relationship between cell-type-specific gene expression and brain-wide connectivity is not well understood. By integrating single-cell resolution spatial transcriptomics and connectomics, we reveal tight gene-connectivity coupling in the cortico-thalamic circuit. To uncover the latent factors associating connectivity with gene expression, we focused on the often-overlooked extrasomatic mRNAs in spatial transcriptomics and identified specific gene expression in axons of the corpus callosum that reflect their cortical origins. Building on these findings, we developed SpaCon, a deep-learning method that flexibly integrates global connectivity with local gene expression. SpaCon employs efficient neighbor-sampling to enable whole-brain analysis while preserving performance. This architecture allows the model to identify functionally relevant three-dimensional domains defined by transcriptome-connectome patterns, even when spatially distant. Validated across diverse datasets and species, SpaCon significantly enhances the prediction of connectivity from gene expression and improves the spatial classification of neuronal subtypes. SpaCon provides a powerful, scalable, and versatile framework for understanding transcriptome-connectome relationships.

## Introduction

The organization and function of the brain arise from the diverse molecular composition across different regions and the complex connections linking them. Understanding how patterns of gene expression relate to brain connectivity is fundamental to deciphering the principles of brain organization ^1,2^. Previous studies have established correlations between regional gene expression profiles and connectivity features. For example, connected regions often share similar transcriptomes ^3–5^, projection targets possess distinct molecular signatures ^6,7^, and connectivity hubs exhibit unique transcriptional profiles ^8–10^. These findings collectively suggest a strong correspondence between the brain’s molecular and connectivity architecture.

Despite these important advances, our understanding of the molecular underpinnings of brain connectivity remains incomplete. First, most prior analyzes primarily used bulk or spatial transcriptomic data without single-cell resolution ^3–9,11–13^. The lack of single-cell resolution prevented the dissection of cell-type-specific contributions to the observed correlations and thus the ability to link molecular identity to circuit architecture. Second, the mechanisms behind these links were often attributed to factors such as coordinated neuronal activity or shared developmental programs ^14,15^, potentially overlooking the direct contribution of extrasomatic mRNAs. Although often discarded as background noise in standard segmentation pipelines, emerging evidence highlights the functional significance of this ‘dark transcriptome’ ^16,17^ and the organized distribution of mRNAs outside cell bodies ^18^. However, the specific contribution of these extrasomatic transcripts to the coupling between gene expression and brain connectivity remains largely unexplored. Neglecting these signals misses the molecular cargo transported within axons—the physical substrate of neural connections ^19–22^. Third, existing computational tools for spatial transcriptomics were designed to integrate transcriptomic data with local features, such as spatial coordinates and histological information ^23–26^, but were inefficient and poorly suited for integrating whole-brain connectivity maps. A lack of analytical frameworks capable of integrating high-resolution spatial transcriptomics with large-scale connectomics has impeded our ability to systematically link molecular profiles to the brain’s global circuit architecture.

Here, we investigate the relationship between gene expression and brain connectivity using the latest spatial transcriptomics ^27–29^ and connectomics datasets ^30,31^. Our analysis reveals a tight, cell-type-specific coupling between gene expression and anatomical connectivity in the cortico-thalamic circuit. By specifically interrogating the spatially resolved transcripts typically unassigned to cell bodies, we provide mechanistic insight revealing that mRNAs within the axon-rich corpus callosum carry transcriptional signatures that reflect their cortical origins. To computationally bridge the gap between the transcriptome and connectome, we introduce SpaCon, a scalable deep-learning framework based on a graph attention autoencoder for multi-modal data fusion. By incorporating a highly efficient neighbor sampling strategy, SpaCon overcomes critical computational barriers to integrate these large-scale, whole-brain multimodal datasets. The resulting fusion of connectomic and transcriptomic data not only enhances the prediction accuracy of connectivity patterns from gene expression, but also improves the classification of neuronal subtypes in cases where spatial transcriptomics alone is insufficient. These results underscore the potential and advantages of SpaCon in integrating multimodal data, providing a powerful method for decoding the principles of brain organization and function.

## Results

### Corticothalamic connectivity is tightly coupled with cell-type-specific gene expression

To investigate the molecular underpinnings of connectivity, we first focused on the well-characterized mouse corticothalamic circuit, an ideal system for this analysis due to its strong projections (**Fig. S1**) and distinct topological patterns **(Fig. S2a)**. Using Allen mouse brain connectivity data ^30^, we first constructed a corticothalamic connectivity matrix, where each thalamus spot was represented by its incoming cortical projection profile. UMAP dimensionality reduction applied to this matrix segregated these spots into eight distinct groups (TH1-TH8), each exhibiting unique connection patterns **(Fig. 1a, S2a-b)**. To determine if these connectivity-based groups possess distinct molecular identities, we analyzed spatially aligned whole-brain spatial transcriptomic data (MERFISH) from mice ^27^, focusing on major thalamic cell types. UMAP analysis of the gene expression profiles for thalamic glutamatergic neurons revealed clusters based on molecular similarity. Notably, when coloring these neurons in the gene-expression UMAP space according to their predetermined connectivity group (**Fig. 1b, S2c**), we observed that neurons from the same connectivity group clustered together, indicating a strong correspondence between corticothalamic connectivity patterns and gene expression profiles in the thalamus. A similar, though less distinct, relationship was observed for thalamic GABAergic neurons, whose connectivity groups also showed some segregation in the gene expression space (**Fig. S2d**). In contrast, the gene expression profiles of glial cells showed no correlation with the connectivity-based groupings (**Fig. S2e**). Thus, the relationship between anatomical connectivity patterns and molecular profiles is a specific feature of thalamic neurons, with the association being strongest in the glutamatergic population.

**Figure 1.**
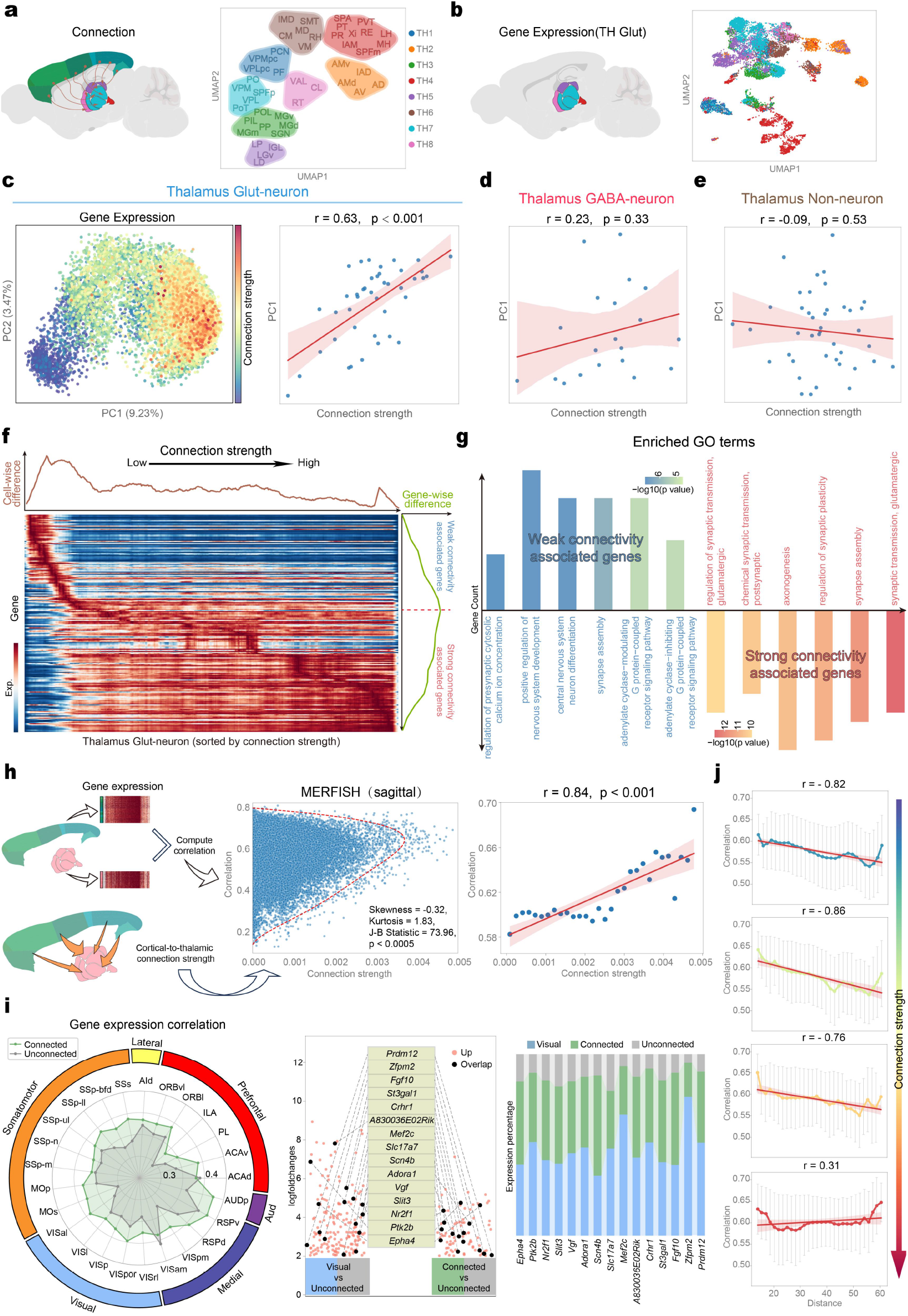
Relationship between corticothalamic gene expression and connectivity. **(a)** Corticothalamic projections and clustering of thalamic nuclei. Based on the connection strength matrices from 43 cortical regions to 44 thalamic nuclei, thalamic nuclei were clustered into 8 distinct categories. Nuclei from different categories receive distinct patterns of cortical projections. *Left*: Schematic of projections. *Right*: UMAP clustering based on projection patterns. **(b)** UMAP clustering of thalamic glutamatergic neurons by gene expression from sagittal MERFISH data, color-coded by corticothalamic projection clusters. Each point in the UMAP plot represents a thalamic glutamatergic neuron. **(c)** Relationship between cortical-to-thalamic connection strength and PC1 of gene expression in thalamic glutamatergic neurons. *Left*: Scatter plot of the first two principal components (PC1, PC2) obtained from PCA on thalamic glutamatergic neuron gene expression data, with each point color-coded by the average cortical-to-thalamic connection strength. The connection strength was defined as the normalized projection volume, following the definition used in the Allen Mouse Brain Connectivity Atlas. *Right*: Scatter plot shows the correlation between PC1 and connection strength. Each point corresponds to a thalamic nucleus; the red line is the linear regression fit across all nuclei, and the shaded region indicates the 95% confidence interval. The *r* denotes the Pearson correlation coefficient, and *p* denotes P-value. **(d)** Similar to (c) *Right*, but based on thalamic GABAergic neurons. **(e)** Similar to (c) *Right*, but based on thalamic non-neuronal cells. **(f)** Gradient of thalamic gene expression along the connection strength from the cortex to the thalamus. Based on the average connection strength, the top 200 genes in the thalamus exhibiting gradient expression with connection strength were identified. The heatmap displays the expression levels of these genes, with thalamic cells sorted by connection strength. The brown line plot in the figure represents the gene expression difference between adjacent cells in the heatmap, and the green line plot represents the expression difference of adjacent genes in the heatmap. According to the position with the largest expression difference (red line), the genes are divided into weak connectivity associated genes and strong connectivity associated genes. **(g)** GO enrichment analysis of weak connectivity associated genes (blue) and strong connectivity associated genes (red). In the bar plot, the height of each bar represents the number of genes enriched for the corresponding GO term, and the bar color indicates the p-value for that term. **(h)** Relationship between cortico-thalamic gene-expression correlation and connection strength. *Left*: Schematic illustration of the relationship between cortico-thalamic gene-expression correlation and connection strength. *Middle*: Scatter plot of connection strength versus gene-expression correlation for cortical-thalamic areas. Each point in the plot represents a cortex-thalamus pair. The red dashed line is the smoothed kernel density estimate of all points. The skewness of the smoothed contour is −0.32, and the kurtosis is 1.83 (normal distribution: skewness = 0, kurtosis = 3). The Jarque-Bera test yields a JB statistic of 73.96 (p < 0.0005), indicating a significant deviation from normality. *Right*: The scatter plot of connection strength versus gene-expression correlation was divided into 30 bins of equal width based on connection strength. The 30 points in the plot represent the median correlation within each bin, ordered by increasing connection strength. A linear regression was fitted to these 30 points; the red line shows the fitted regression line, and the shaded area indicates the 95% confidence interval. **(i)** *Left*: Gene-expression correlation between cortical regions and connected (green) vs. unconnected (gray) thalamic areas. *Middle*: DEGs between the visual cortex and connected vs. unconnected thalamic nucleus. The red points on the left represent genes upregulated in unconnected thalamic regions compared to visual cortical regions, while the red points on the right represent genes upregulated in thalamic regions connected to the visual cortex compared to unconnected thalamic regions. Black points denote genes common to both sets, with their names listed in the center. *Right*: Expression levels of DEGs in the visual cortex (blue), connected (green), and unconnected (gray) thalamic regions. **(j)** Relationship among cortico-thalamic gene-expression correlation, connection strength, and spatial distance. All cortico-thalamic point pairs were divided into four intervals based on connection strength; each panel shows the relationship between gene-expression correlation and spatial distance for point pairs within each interval, with error bars representing one standard deviation, the red line indicating the fitted regression line, and the shaded area representing the 95% confidence interval.

To quantify the relationship between gene expression and connection strength across cell types, we performed Principal Component Analysis (PCA) on the gene expression profiles of thalamic glutamatergic neurons, GABAergic neurons, and non-neuronal cells, separately. For glutamatergic neurons, visualization of the first two principal components (PCs) showed a clear gradient corresponding to the strength of corticothalamic connections, particularly along PC1 (**Fig. 1c, S3a**). This relationship was confirmed by a significant positive correlation between the average gene expression PC1 values and connection strengths within defined thalamic regions (r = 0.63, p < 0.001; **Fig. 1c**), demonstrating that the dominant axis of transcriptional variation in thalamic glutamatergic neurons is closely linked to the strength of their cortical input. Conversely, analogous analyzes for thalamic GABAergic neurons (**Fig. 1d, S3b**) and non-neuronal cells (**Fig. 1e, S3c**) showed substantially weaker and non-significant correlations. These results provide quantitative evidence that the coupling between corticothalamic connectivity patterns and gene expression profiles is largely specific to the glutamatergic neurons in the thalamus.

To characterize the molecular correlates of corticothalamic connectivity, we used scVelo method ^32^ to identify genes in thalamic glutamatergic neurons whose expression correlated with connection strength. Visualization of the top 200 correlated genes revealed distinct expression modules clearly stratified by connection strength, with many genes exhibiting sharp, threshold-like changes in expression rather than gradual shifts (**Fig. 1f**). Functionally, these gene sets were highly divergent. Gene Ontology (GO) analysis ^33^ showed that genes associated with strong connectivity were enriched for synaptic transmission and plasticity, while those associated with weak connectivity were enriched for nervous system development, synapse assembly, and specific signaling pathways (**Fig. 1g**). These results suggest that cortical input strength is coupled to functionally distinct transcriptional features in thalamic glutamatergic neurons.

We next investigated whether the relationship between connectivity and gene expression extends to transcriptional correlation between connected cortical and thalamic regions. Analyzing the relationship between the pairwise transcriptional correlation of cortical and thalamic spots and their corresponding anatomical connection strength revealed a skewed distribution (**Fig. 1h**). This distribution indicates that while weak connections show variable correlation, strong connections are almost exclusively associated with high transcriptional correlation. Grouping the data into 30 bins by connection strength further clarified this relationship, revealing a strong, significant positive correlation (r = 0.84, p < 0.001) between connection strength and the average gene-expression correlation (**Fig. 1h, S3e**). This pattern was consistent across the corticothalamic system, with connected areas exhibiting significantly higher expression correlation than unconnected areas for all 25 cortical regions examined (**Figs. 1i, S4**). Differential expression analysis identified genes (DEGs) highly expressed in connected regions, many of which are involved in synapse functions and ion channel activity (**Fig. S5**), highlighting their roles in brain connectivity.

Given that gene-expression correlation may decay with increasing spatial distance between regions ^2,8,14^, we investigated whether strong connectivity could counteract this effect. We divided the cortico-thalamic connections into four quartiles based on their strength and examined the relationship between spatial distance and gene-expression correlation within each quartile (**Fig. 1j**). For the three quartiles of weaker connections, correlation decayed significantly with increasing distance between cortical and thalamic spots. In contrast, for the quartile of the strongest connections, transcriptional correlation remained uniformly high and showed no significant correlation with distance (r = 0.31). These results demonstrate that strong anatomical connectivity is associated with a high degree of transcriptional correlation between brain regions, regardless of spatial distance.

### Extrasomatic axonal transcripts reveal signatures of their cortical origins

Our findings in the cortico-thalamic system strongly imply a physical mechanism that bridges the transcriptomes of connected regions. In addition to known factors like mutual developmental influences and coordinated activity ^14,15^, we hypothesized this link is also contributed by the direct transport of mRNA along axons. Validating this hypothesis requires detecting transcripts located physically distant from the cell soma—signals that are typically classified as ‘extrasomatic’ and discarded in standard segmentation pipelines. However, emerging studies have demonstrated that these unassigned transcripts are not random noise but possess functional relevance and organized distribution ^17,18^. Leveraging this insight, we sought to isolate axonal mRNA signatures by focusing on the corpus callosum. Unlike gray matter regions where distinguishing axonal transcripts from somatic overflow is challenging, the corpus callosum is a major white matter tract comprised of long-range cortical axons and glial cells, but devoid of neuronal cell bodies. This provides an ideal structure to isolate and identify these potential axonal mRNA signatures.

We first examined whether the axon-enriched extracellular compartment of the corpus callosum possesses distinct gene expression patterns compared to the surrounding intracellular (cell body-enriched) glial compartments. Differential expression analysis based on spatial transcriptomic segmentation uncovered numerous DEGs between different compartments (**Fig. 2a-b, S6**). While intracellular-upregulated genes were most enriched for markers of resident glia like oligodendrocytes and precursors (OPC-Oligo), extracellular-upregulated genes displayed the strongest enrichment for glutamatergic neuron markers. For example, spatial expression maps demonstrated higher relative abundance of the glutamatergic marker *Slc17a7* ^27^, the axon-associated gene *Fgf13* ^34^, and the layer 2/3 cortical neuron marker *Calb1* ^35^ within the extracellular compartment (**Fig. 2b, S7a**). To ensure this enrichment was not an artifact of the substantially larger area of the extracellular compartment, we normalized mRNA counts by area and confirmed the findings remained significant (**Fig. S7b**). The differential expression patterns were highly reproducible, showing significant overlap in the top 100 differentially expressed genes between coronal and sagittal sections analysis (**Fig. 2c**) and a strong correlation in their respective log_2_ fold-change values across all DEGs (r = 0.95, p < 0.001; **Fig. 2d**). GO enrichment analysis indicated that extracellular-enriched genes are predominantly associated with axonal structures and synaptic processes (e.g., ‘distal axon’, ‘axon terminus’, ‘synaptic vesicle’), whereas intracellular-enriched genes are linked to glial functions and extracellular matrix components (e.g., ‘collagen-containing extracellular matrix’, ‘receptor complex’, ‘adherens junction’) (**Fig. 2e**). Furthermore, this compartment exhibited elevated expression of marker genes characteristic of cortical layers 2/3 and 5 compared to other layers (**Fig. 2f, S8a-b**), consistent with previous findings that interhemispheric callosal projections originate predominantly from these layers ^31,36^.

**Figure 2.**
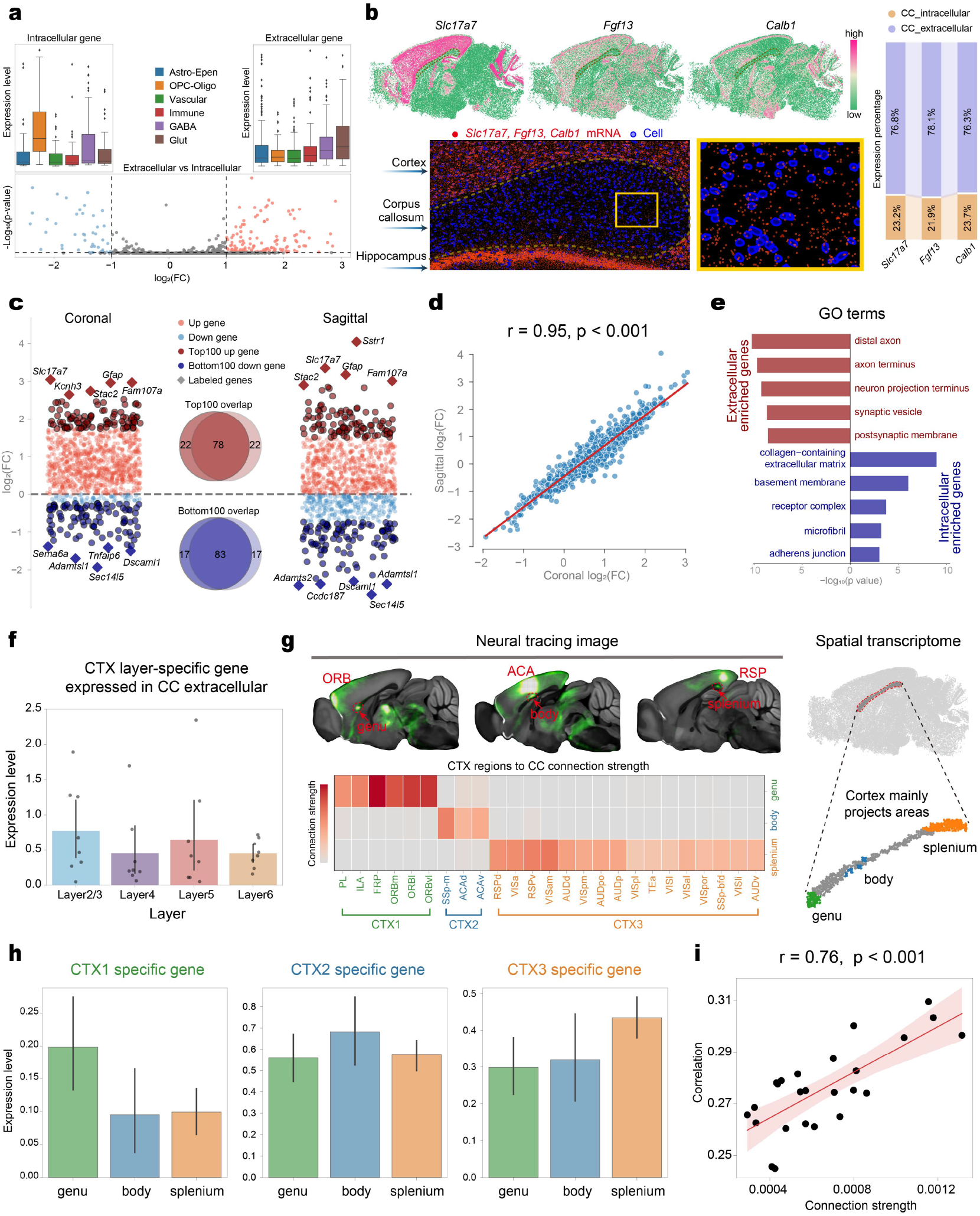
Analysis of connectivity and gene expression between cortical and corpus callosum compartments. **(a)** Differential expression analysis of intracellular and extracellular compartments of the corpus callosum. *Bottom*: Differentially expressed genes (DEGs) between intracellular and extracellular compartments. *Top*: Average expression of intracellular and extracellular DEGs across different cell types. Astro, astrocytes; Epen, ependymal cells; OPC, oligodendrocyte precursor cells; Oligo, oligodendrocytes; Vascular, vascular cells; Immune, immune cells; GABA, GABAergic neurons; Glut, glutamatergic neurons. Box plots display the median, 25th (Q1) and 75th (Q3) percentiles, with whiskers extending to 1.5× the interquartile range (IQR); outliers beyond this range are marked individually. ***(b)*** *Left*: Sagittal sections showing gene expression of *Slc17a7* (glutamatergic neuron marker), *Fgf13* (axon-related gene), and *Calb1* (layer 2/3-specific neuron marker). Enlarged panels highlight mRNA localization (red) and nuclei (blue). *Right*: Gene expression ratios in intracellular and extracellular compartments of the corpus callosum (CC). **(c)** Differential expression analysis between intracellular and extracellular compartments of the corpus callosum in both MERFISH coronal and sagittal datasets. Dot plots show the distribution of log fold changes of all genes. In each plot, dark red dots represent the top 100 upregulated genes, and dark blue dots represent the top 100 downregulated genes. Diamond-shaped points with gene names highlight the top five most upregulated and downregulated genes in each dataset. Venn diagrams show overlaps of the top 100 upregulated and downregulated genes across datasets. **(d)** Correlation of log fold changes across all genes between coronal and sagittal datasets. Each dot represents one gene. The x-axis of each point indicates the log fold change (extracellular vs intracellular) of the corresponding gene in the coronal dataset, and the y-axis indicates the log fold change in the sagittal dataset. The red line indicates the fitted regression line. The *r* denotes the Pearson correlation coefficient, and *p* denotes P-value. **(e)** GO enrichment analysis of the overlapping genes in (c) (78 upregulated, 83 downregulated in extracellular vs. intracellular compartments). Red bars represent upregulated genes; blue bars represent downregulated genes. **(f)** Bar plot showing extracellular expression of layer-specific marker genes (8 genes per layer) in the corpus callosum region. Each point represents the expression level of a layer-specific gene in the extracellular compartments of the corpus callosum. Error bars represent the 95% confidence interval. **(g)** Cross-hemispheric projections via different parts of the corpus callosum. *Left top*: Neuronal tracing images from the Allen Brain Atlas showing injections in different cortical regions (ORB, orbital area; ACA, anterior cingulate area; RSP, retrosplenial area), source: https://connectivity.brain-map.org/. *Left bottom*: Connection strength from 25 cortical regions to 3 corpus callosum regions. Based on each cortical area’s primary projection to regions of the corpus callosum, the cortex was classified into three parts—CTX1, CTX2, and CTX3—which primarily project to the genu, body, and splenium of the corpus callosum, respectively. *Right*: Spatial distributions of the corresponding corpus callosum regions in the spatial transcriptomics data: genu (green), body (blue), splenium (orange). **(h)** Expression of cortical region-specific genes (DEGs in CTX1, CTX2, and CTX3) in extracellular compartments of the corpus callosum. The cortex and corpus callosum were divided into three regions each according to cortico-callosal projections (Fig. 2g). Bar plots show expression levels of the top 5 DEGs from each cortical region (CTX1, CTX2, or CTX3) within extracellular compartments of the corpus callosum. Error bars denote 95% confidence intervals. **(i)** Association between connection strength and gene-expression correlation across cortical-callosal regions. Each data point corresponds to a cortical-callosal region pair (25 cortical regions×3 callosal regions). The x-axis shows connection strength between paired regions, while the y-axis shows their gene-expression correlation. In the figure, r represents the Pearson correlation coefficient of all points, and p represents the corresponding p-value.

To determine whether the axonal molecular signature could reflect cortical origins, we integrated neural tracing data with our spatial transcriptomic maps. We focused on three distinct corticocallosal subdivisions, where cortical regions CTX1, CTX2, and CTX3 project preferentially through the genu, body, and splenium, respectively (**Fig. 2g, S8c-e**). This revealed a remarkable topographic mapping: genes enriched in each cortical group were also enriched in the extracellular compartment of their corresponding callosal target, compared to non-target subdivisions (**Fig. 2h, S8f**). Moreover, the strength of corticocallosal connections correlated strongly with the transcriptional correlation between projecting cortical regions and their target axonal compartments (**Fig. 2i, S8g**). Taken together, these findings demonstrate that the extracellular (axon-enriched) compartment of the corpus callosum carries a specific repertoire of mRNAs that reports their anatomical source and connection strength. This supports the hypothesis that axonal mRNA transport may contribute to the observed transcriptomic correlation between anatomically connected brain regions.

### Tunable and efficient integration of local gene expression and global connectivity via SpaCon

Our findings demonstrate that anatomical connectivity and gene expression are closely related within brain circuits (**Figs. 1-2**). This tight relationship necessitates analytical approaches that can effectively integrate these distinct data modalities for a more comprehensive view of brain organization. However, integrating these data types presents a significant challenge. Spatial transcriptomics provides rich local molecular features, while connectomics describes the global network architecture. Current analytical tools of spatial transcriptomics were primarily optimized for integrating spatially adjacent multi-modal features and were therefore not designed to handle the global connectivity information ^30^. Furthermore, the scale and complexity of global connectivity information introduce significant computational challenges when attempting to integrate multi-modal data across entire brains. To address these challenges, we introduce SpaCon (Spatial Transcriptomics and Connectomics Integration), a deep learning method that effectively integrates whole-brain connectome information with spatial transcriptomics data.

SpaCon employs a graph neural network (GNN) framework, built upon a graph attention autoencoder architecture ^37^, to achieve the integration of multi-modal data (**Fig. 3a**). We first represent these two data modalities (connectome and transcriptome) within a unified graph structure that serves as the model’s input. This structure comprises two conceptual graph types: a Spatial Graph and a Connection Graph. In both graphs, nodes represent spatial transcriptomic spots (or cells) featuring their gene expression profiles. Edges in the Spatial Graph are defined by local spatial proximity between nodes, thus encoding local spatial information, whereas edges in the Connection Graph represent connectivity features between nodes, thus encoding global connectivity patterns. The SpaCon model consists of two specialized encoders (Spa_encoder and Con_encoder) and a decoder. The Spa_encoder and Con_encoder process the Spatial and Connection Graphs, respectively, learning latent representations for each modality. These representations are subsequently merged via a feature fusion module to generate a unified latent embedding. Finally, this embedding is passed to the decoder to reconstruct the original gene expression matrix, and the entire network is trained by minimizing the reconstruction loss, thereby optimizing the integrated representation.

**Figure 3.**
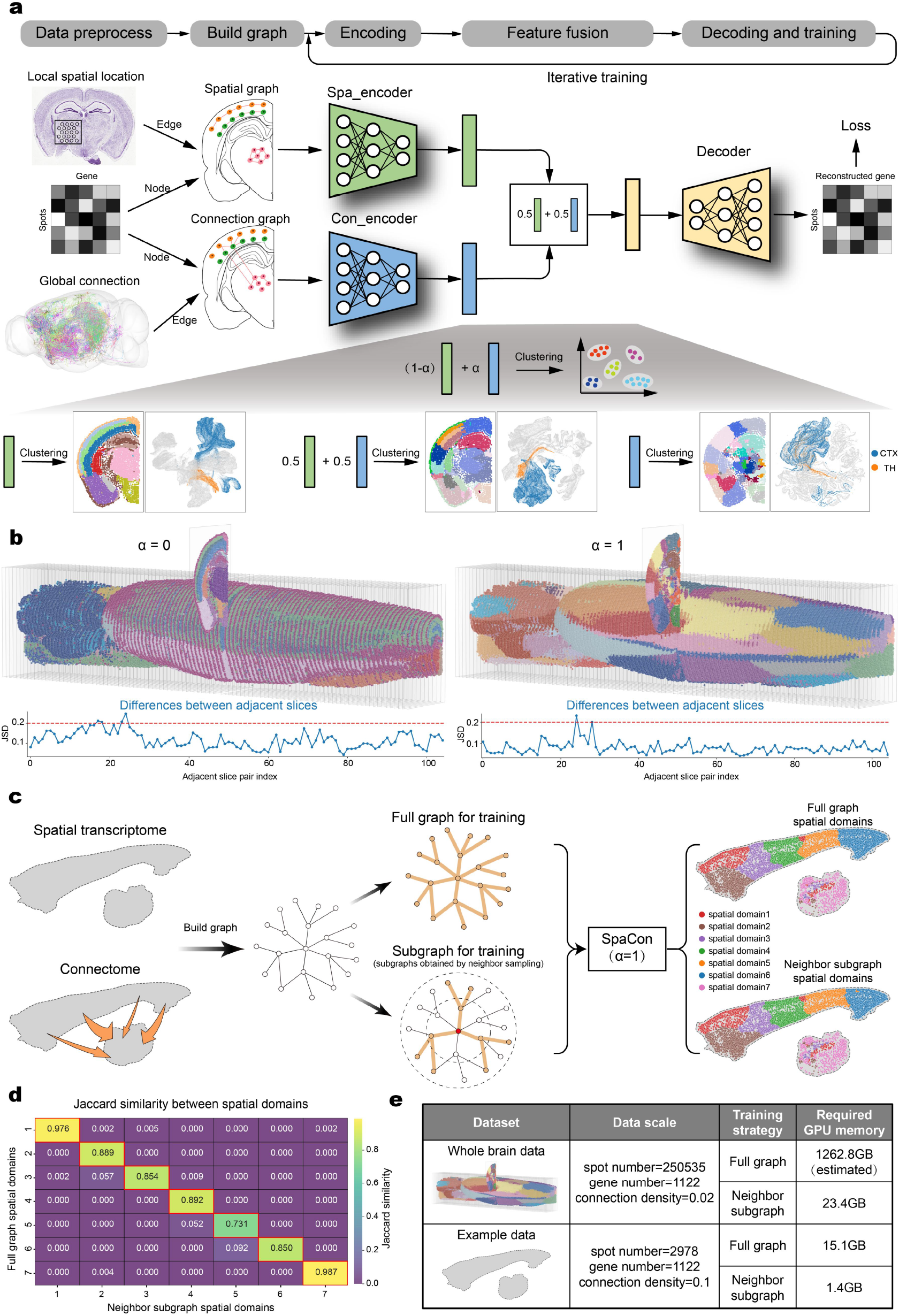
SpaCon workflow. **(a)** SpaCon workflow. Gene expression data (nodes) were integrated with spatial information (edges) and connectivity information (edges) to construct a spatial graph and a connectivity graph, respectively. These graphs were processed by two separate encoders, and their output features were fused by addition. The fused features were passed through a decoder to reconstruct gene expression for loss computation. After iterative training, the encoder outputs were combined using a tunable parameter α to balance spatial and connectivity contributions, and a clustering algorithm was applied to identify spatial domains. Below we show the clustering results obtained under different parameters and the corresponding UMAP plots. In the coronal slice, different colors indicate different spatial domains. In the UMAP plots, blue points represent cortex (CTX), orange points represent thalamus (TH) and gray points represent other regions. **(b)** Three-dimensional clustering results obtained by SpaCon. *Top*: Clustering results for a whole mouse brain slice with α set to 0 and 1. Different colors represent distinct spatial domains identified by SpaCon. All coronal slices are arranged along the rostro-caudal axis. *Bottom*: Line plots depict Jensen-Shannon divergence (JSD) between adjacent slices, where the red dashed line (JSD=0.2) indicates the empirical threshold for negligible batch effects (JSD<0.2 suggests smaller inter-section variability). **(c)** Schematic comparison of full-graph training versus subgraph sampling (subgraphs are obtained by neighbor sampling). For graphs constructed from spatial transcriptomics and connectomics data, clustering was performed using neighbor-based subgraph sampling and full-graph training, respectively, yielding the corresponding clustering results. **(d)** Heatmap showing pairwise Jaccard similarity coefficients between spatial domains identified by full graph clustering (y-axis) and neighbor subgraph clustering (x-axis). Color scale represents Jaccard similarity index values (intersection over union), with higher values indicating greater clustering consistency. **(e)** Comparison of GPU memory usage. The table presents GPU memory usage for both whole-brain data and example data (consisting of cortical-thalamic regions from two slices of the whole-brain dataset) under two training strategies. The ‘spot number’ and ‘gene number’ denote the counts of spots and genes in each dataset; ‘connection density’ is the proportion of nonzero entries in the adjacency matrix; ‘full graph’ refers to model training on the full graph; ‘neighbor graph’ refers to mini-batch training after neighbor sampling. Since full graph training cannot be applied to the whole-brain dataset, the GPU memory usage is an estimated value.

A key component of our model is the feature fusion mechanism, governed by a tunable hyperparameter, α, which weights the relative contribution of the Spa_encoder and Con_encoder outputs to the final embedding (**Fig. 3a**). This parameter controls the balance between local spatial information and global connectivity information. Lower α values prioritize local spatial context, leading to clustering results with strong spatial coherence. Conversely, higher α values emphasize global connectivity patterns, enabling the identification of potentially spatially discontinuous regions linked by connectivity information. During the model training phase, we recommend setting the α to 0.5, so that the model learns the relationship between the two modalities by giving equal weight. After training is complete, α can be tuned to adjust the final output, allowing users to flexibly generate clustering results that emphasize either local spatial patterns (α close to 0) or global connectivity architecture (α close to 1), depending on the specific application (e.g., α=0 vs. α=1; **Fig. 3b, S9**). Analysis of inter-slice consistency using Jensen-Shannon Divergence (JSD) confirmed that α provides tunable control over the stability of 3D domain boundaries across adjacent slices (**Fig. 3b**). This demonstrates SpaCon’s ability to overcome potential batch effects inherent in multi-slice spatial transcriptomic data (**Fig. S10**), thereby ensuring robust identification of 3D domains and enhancing the reproducibility of biologically meaningful results.

A major computational bottleneck in integrating whole-brain multi-modal data is their large size, which often exceeds the memory capacity of standard GPUs. To overcome this, we implemented a connectivity-based neighbor sampling strategy within the SpaCon framework (**Fig. 3c**). Our method avoids processing the full graph by operating on small, information-rich subgraphs during each training step. A subgraph is built around a central ‘anchor’ node and is populated by its most relevant neighbors—those with strong connections and close network proximity. This design ensures that each training step is focused on a computationally manageable yet biologically meaningful portion of the network, capturing both direct connections and local circuit topology. This sampling strategy proved highly effective, accurately replicating the results of full-graph training while reducing the computational load. Quantitative comparison confirmed that, compared with the random sampling method (**Fig. S11**), the spatial domains identified via neighbor sampling were nearly identical to those from the full-graph method (**Fig. 3c**), with exceptionally high pairwise Jaccard similarities between corresponding domains ranging from 0.731 to 0.987 (**Fig. 3d**). Crucially, this approach resolved the computational bottleneck associated with large datasets by reducing GPU memory usage by over 90% compared to the full-graph strategy (**Fig. 3e**), enabling SpaCon to scale effectively to whole-brain analyzes.

### SpaCon reveals spatial domains with integrated molecular and connectivity profiles

We applied SpaCon to the mouse cortico-thalamic system, training the model to integrate anatomical connectivity from Allen mouse brain connectivity data ^30^ and gene expression from Stereo-seq or MERFISH spatial transcriptomics data ^27,29^ with a balanced weight (α=0.5). Subsequent clustering, configured to prioritize long-range connectivity (α=1), robustly grouped specific cortical areas with their principal thalamic targets into unified 3D spatial domains (**Fig. 4a, S12a**). For example, it grouped the retrosplenial/motor cortex (RSP/MO) with the lateral dorsal nucleus (LD), the somatosensory cortex (SS) with the ventral posteromedial nucleus (VPM), and the auditory cortex (AUD) with the medial geniculate nucleus (MG). The co-clustering patterns were highly reproducible across independent Stereo-seq and MERFISH datasets (Pearson’s r = 0.836, p < 0.0001) and correlated significantly with anatomical connectivity patterns (Pearson’s r = 0.739, p < 0.0001), confirming their biological relevance (**Fig. S12b**). Notably, these spatial domains reflect an integrated molecular and connectional identity. Connected cortical and thalamic spots within the same SpaCon domain showed significantly higher gene-expression correlation than those in different domains (**Fig. 4b, S12c**). Furthermore, this integration was demonstrated by the finding that the first principal component of gene expression from a specific thalamic domain more accurately predicts its connection strength to its matched cortical partner than to any other cortical region (**Fig. 4c-d, S13a**). These results indicate that SpaCon learns a unified representation in which molecular profiles and connectivity are mutually predictive.

**Figure 4.**
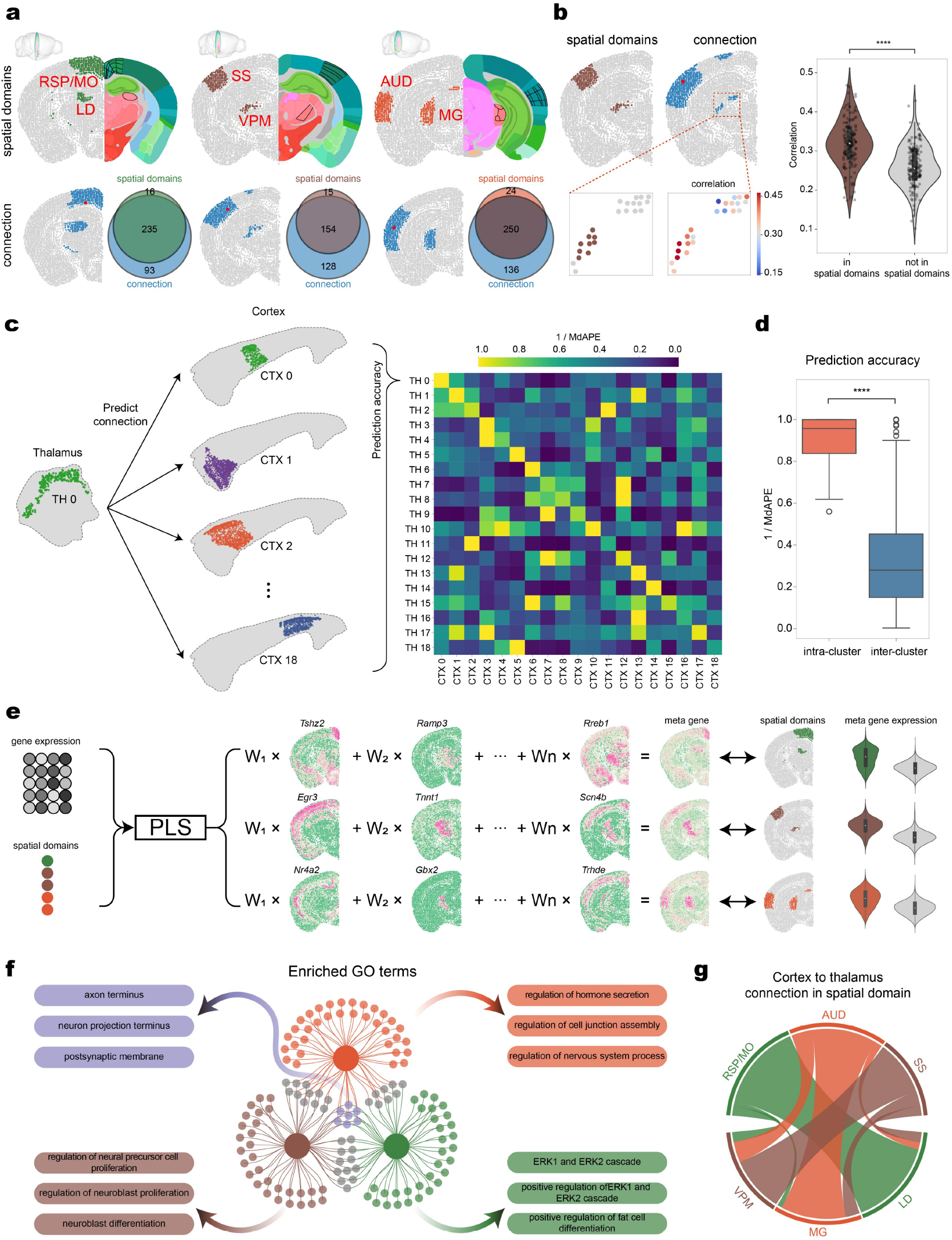
SpaCon integrated clustering of the cortex and thalamus. **(a)** Clustering results of cortex and thalamus when α = 1. *Top*: Spatial domains involving cortical areas and thalamic nuclei. The 3D view in the upper left corner indicates the position of the corresponding slice. In each slice, the left half shows the clustering results of cortex and thalamus, and the right half shows the corresponding Allen atlas, with regions outlined in black denoting the brain areas matching the clustering results. RSP, retrosplenial cortex; MO, motor cortex; LD, lateral dorsal nucleus; SS, somatosensory cortex; VPM, ventral posteromedial nucleus; AUD, auditory cortex; MG, medial geniculate nucleus. *Bottom*: Corresponding connectivity maps (*left*: injection sites shown in red spots) and the overlap between the connectivity maps and the spatial domains (*right*). The numbers in the Venn diagram represent the number of spots. ***(b)*** *Left*: Two coronal slices showing SpaCon spatial domains (left) and the corresponding connectivity graph (right). Red dots mark injection sites sampled from the spatial domain. The thalamic region in the connectivity graph is enlarged: in the left inset, brown dots represent points overlapping the spatial domain and gray dots represent points outside the spatial domain; in the right inset, dot colors indicate gene-expression correlation with the injection sites. *Right*: For each cortical point in the spatial domain used as an injection site, a corresponding connectivity graph is generated. Violin plots summarize the gene-expression correlations for points overlapping (‘in spatial domain’) and not overlapping (‘not in spatial domain’) the spatial domain in each connectivity graph. **(c)** Predictive relationships of corticothalamic spatial domains based on SpaCon clustering. *Left*: Prediction schematic showing how PC1 of gene expression in each thalamic domain predicts the connection strength from each cortical domain. Cortical-thalamic pairs with the same numeric label belong to the same spatial domain (e.g. TH1 and CTX1). *Right*: Prediction accuracy (1 / MdAPE) for each thalamic domain predicting each cortical domain. **(d)** Box plots comparing intra-domain (orange) and inter-domain (blue) prediction accuracy. Intra-domain refers to cortical-thalamic spatial domains with matching numeric labels in (c), while inter-domain refers to mismatched domains. **(e)** Metagenes identified for each spatial domain via PLS. For the spatial domains obtained by SpaCon, PLS fitting assigns each gene a weight coefficient representing its association with the spatial domain. For each spatial domain, we selected the 60 genes with the highest absolute weight values and linearly combined them into a ‘metagene’. Right violin plots display distinct metagene expression profiles within and outside the spatial domains. **(f)** Overlap and unique gene sets for metagenes in three spatial domains, along with the corresponding GO terms. The central Venn diagram shows the intersection of metagenes (the top 60 genes by absolute PLS weight) among the three spatial domains in panel (a). Arrows emanating from the overlapping regions indicate the principal GO enrichment pathways associated with those genes. **(g)** Connectivity patterns among different cortical areas and thalamic nuclei. The chord diagram shows the mutual connection strengths among the three spatial domains in panel (a), with ribbon thickness indicating the magnitude of connectivity. Cortical source regions and thalamic target regions belonging to the same spatial domain are displayed in the same color.

To reveal the molecular basis of these domains, we employed partial least squares (PLS) regression ^38^ to derive ‘metagenes’, linear combinations of genes whose expression optimally distinguished each spatial domain. These metagenes captured the unique transcriptional identity of each cluster, exhibiting significantly elevated expression within their corresponding domains (**Fig. 4e, S13b**). GO analysis of these metagenes revealed a molecular organization defined by both domain-specific gene sets and gene sets shared across domains (**Fig. 4f, S13c**). Shared genes were enriched for fundamental processes of neural connectivity, such as axon guidance, neuron projection, and postsynaptic membrane organization, reflecting the shared structural basis of cortico-thalamic circuits. In contrast, the domain-specific gene sets were enriched for distinct biological pathways (e.g., hormone secretion, neuroblast proliferation, ERK cascade regulation), highlighting specialized molecular functions associated with specific cortico-thalamic subsystems. This complex pattern of unique and shared molecular components aligned with the anatomical projection patterns: each cortical region exhibited a unique, dominant connection with its primary thalamic target, while also participating in a shared network of less dominant connections to other thalamic nuclei (**Fig. 4g**). This alignment of molecular and anatomical patterns underscores SpaCon’s ability to resolve functionally relevant subdivisions based on their integrated identities.

### SpaCon enhances cell-type classification in spatial transcriptomics by integrating connectivity

Having demonstrated SpaCon’s ability to define large-scale domains, we next investigated whether integrating connectomics could improve the spatial classification of cell subtypes from spatial transcriptomics data. The classification is sometimes challenging for certain neuronal subtypes, which can be identified by high-depth scRNA-seq but may be unresolved by spatial transcriptomics alone due to its limited gene-detection efficiency ^39^. We hypothesized that the distinct anatomical projection patterns could provide complementary information to improve the classification.

To test this, we focused on layer 6 corticothalamic (CT) neurons, whose subtypes have distinct anatomical projections but are difficult to distinguish using spatial transcriptomics alone. SpaCon was first trained with a balanced hyperparameter (α=0.5), allowing it to learn the underlying relationships between anatomical connectivity ^30^ and spatial transcriptomic features ^27^. For the final classification, we set the hyperparameter to zero (α=0), thereby restricting the model to using only its learned spatial-transcriptomic features to ensure a direct comparison with other methods. Using ground-truth labels previously derived from integrated spatial transcriptomic and snRNA-seq data ^27^ (**Fig. 5a**), we benchmarked SpaCon against four state-of-the-art methods that rely on transcriptomic information, including SpaGCN ^23^, STAGATE ^24^, SEDR ^26^, and SpaceFlow ^25^. Notably, SpaCon robustly identified the spatial patterns of CT neuron subtypes, significantly outperforming all competing methods. The spatial domains defined by SpaCon (**Fig. 5b**) aligned more closely with the ground-truth annotations than those generated by the other methods (**Fig. 5c**). This superior performance was also validated quantitatively across four standard clustering metrics (ARI, NMI, V-measure, and F1 score), where SpaCon consistently achieved the highest scores (**Fig. 5d**).

**Figure 5.**
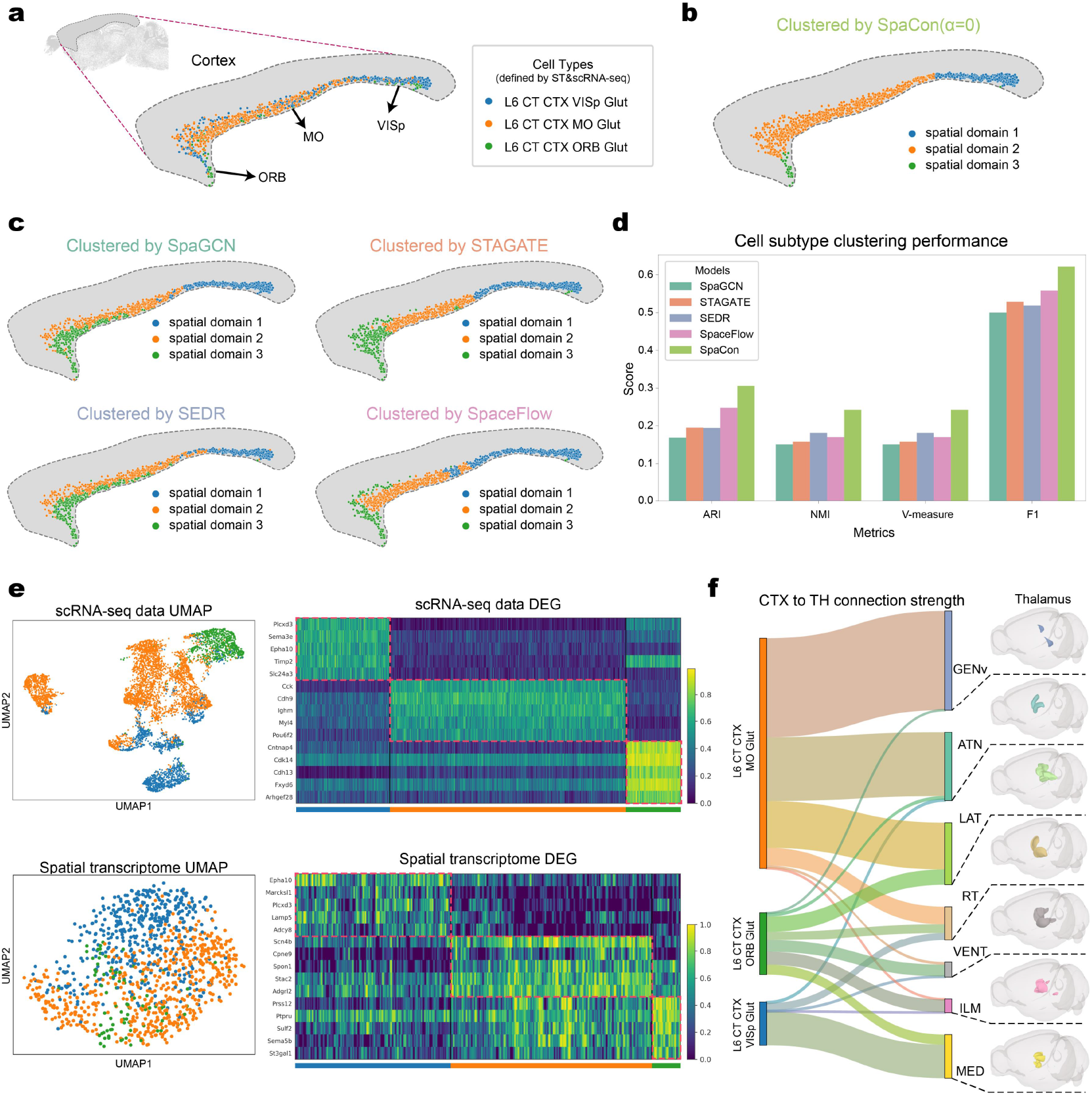
Comparison of SpaCon in classifying neuronal subtypes. **(a)** Cell types defined by the spatial transcriptomics and scRNA-seq data. Cell types in spatial transcriptomics data inferred via mapping from scRNA-seq data of layer 6 corticothalamic (CT) neurons. L6 CT CTX VISp Glut: Corticothalamic glutamatergic neuron subtype located in layer 6 of cortical VISp. L6 CT CTX MO Glut: Corticothalamic glutamatergic neuron subtype located in layer 6 of cortical MO. L6 CT CTX ORB Glut: Corticothalamic glutamatergic neuron subtype located in layer 6 of cortical ORB. **(b)** Clustering results obtained by SpaCon (α=0) based on spatial transcriptome and connectome data. **(c)** Clustering results obtained by SpaGCN, STAGATE, SEDR, SpaceFlow based on spatial transcriptome data. **(d)** Consistency evaluation between clustering results of each model and reference cell types. The bar plot shows the clustering results of each model in the four metrics: ARI (adjusted rand index), NMI (normalized mutual information), V-measure and F1 (F1 Score). **(e)** UMAP embeddings and heatmaps of differentially expressed genes for three CT neuronal subtypes in both scRNA-seq and spatial transcriptomics datasets. Cell type colors match those in (a). The heatmap displays the top five differentially expressed genes for each subtype. **(f)** Connection strength from the three CT neuron subtypes to different thalamic regions. In the Sankey diagram, the width of the connections indicates connection strength. The left side represents cell types, while the right side shows thalamic regions with corresponding 3D spatial maps.

To understand the basis for SpaCon’s performance, we examined the discriminatory power of different data modalities. In the scRNA-seq data ^27^, the three CT subtypes were transcriptionally distinct, forming well-separated clusters and exhibiting clear blocks of DEGs. In contrast, due to the limited gene detection, these same subtypes appeared intermingled in the UMAP plot of spatial transcriptomic data ^27^ with a less defined DEG heatmap (**Fig. 5e, S14a**). However, different transcriptomic-defined subtypes possessed highly specific and segregated projection patterns to distinct thalamic nuclei, providing effective connectivity information for classification (**Fig. 5f, S14b**). SpaCon leveraged these connectivity profiles during training to fuse gene expression patterns linked to connectivity features. This approach enabled the accurate differentiation of CT subtypes and demonstrated that global connectivity provides crucial information where local spatial transcriptomics alone is insufficient.

### Broad applicability of SpaCon across diverse data modalities and species

To demonstrate the versatility of SpaCon, we extended our analyzes beyond anatomical connectomes to integrate spatial transcriptomics with functional connectivity data from different imaging modalities and species.

We first applied SpaCon to the mouse dorsal cortex, integrating spatial transcriptomics ^27^ with large-scale functional connectivity maps derived from wide-field calcium imaging ^36^. After training the model with a balanced integration parameter (α=0.5), we performed the final clustering by prioritizing functional connectivity (α=1). This approach identified several spatial domains that encompassed functionally linked but spatially discontinuous regions. To define the molecular signatures of these domains, we used PLS regression to identify ‘metagenes’ whose composite expression profiles effectively distinguished each cluster (**Fig. 6a, S15a**). GO enrichment analysis revealed these domains were linked to distinct biological processes. For instance, long-range domains (e.g., domain 1, 2, and 3) showed metagene enrichment for ion channel functions, potentially reflecting neuronal communication, whereas a more local domain (e.g., domain 4) was associated with extracellular matrix organization and neurogenesis (**Fig. 6b**). This molecular divergence was also reflected in the functional connectivity profiles of the domains. While all identified domains showed high intra-domain functional connectivity, the long-range domains (e.g., domain 1-3) also maintained strong inter-domain connectivity with each other—a feature absent in the more functionally-isolated local domain 4 (**Fig. 6c**). Thus, SpaCon-defined spatial domains represent cohesive units of brain organization, each characterized by a unique profile of molecular identity and functional connectivity patterns.

**Figure 6.**
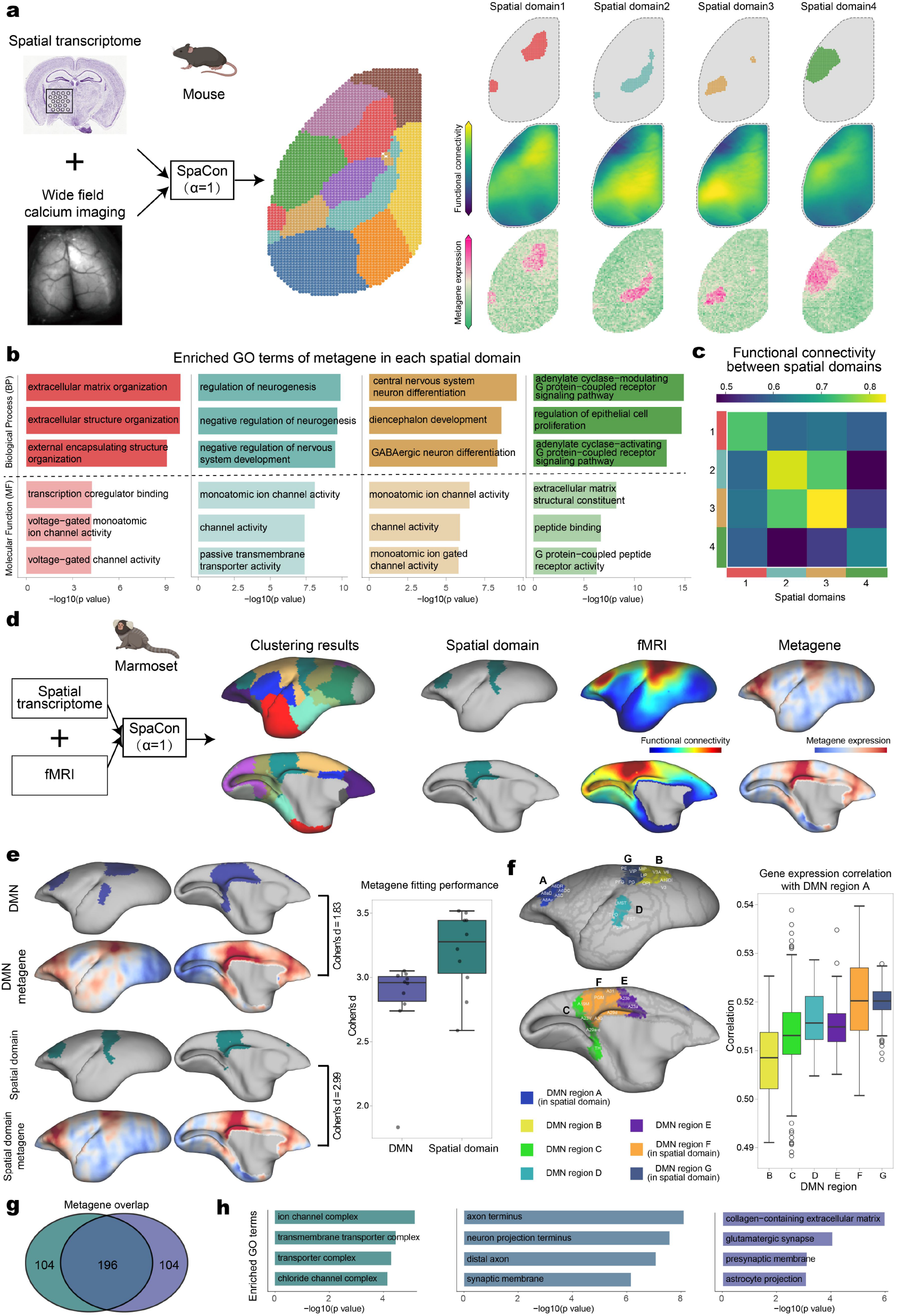
SpaCon enables integration of spatial transcriptomics with wide-field calcium imaging data and fMRI. **(a)** Integrated clustering results of spatial transcriptomics and wide-field calcium imaging data. *Left:* Clustering results of cortical regions. *Right*: First row shows the four spatial domains within the cortex. Second row depicts the functional connection strength of the spots from each domain across the entire cortical region. Third row displays the cortical expression of a set of metagenes identified by PLS within each spatial domain. **(b)** Enriched GO terms of metagene in each spatial domain. GO enrichment analysis was performed on the top 300 metagenes from each spatial domain. GO terms were classified into Molecular Function (MF) and Biological Process (BP) categories, with three terms selected from each category. **(c)** The functional connection strength between the spatial domains in (a). **(d)** SpaCon applied to marmoset spatial transcriptome and fMRI data. *Left:* Clustering results of cortical regions. *Right*: First row shows the two spatial domains within the cortex. Second row depicts the functional connection strength of the spots from each domain across the entire cortical region. Third row displays the cortical expression of a set of metagenes identified by PLS within each spatial domain. **(e)** Comparison of metagene fitting performance between DMN and SpaCon spatial domain. *Left*: DMN and SpaCon spatial domain and their corresponding metagene expressions. Cohen’s d quantifies the metagene expression difference inside vs. outside the domain, higher values indicate a better fit. *Right*: Fitting performance of different numbers of metagenes to the DMN and spatial domains. Each point represents a different number of metagenes, ranging from 100 to 1000 genes in steps of 100 (10 points per group). ***(f)*** *Left*: Based on the overlap between the DMN and SpaCon spatial domains, the DMN was divided into seven regions (A-G), with regions A, F, and G belonging to the same spatial domain. *Right*: The boxplot shows the gene-expression correlation between each spot in DMN regions B-G and region A. **(g)** Overlap of the top 300 metagenes fitted using PLS in the spatial domain and the DMN. **(h)** GO enrichment analysis results of shared and domain-specific metagenes in the spatial domain and the DMN. *Left*: GO terms of SpaCon domain-specific genes. *Middle*: GO terms of shared genes. *Right*: GO terms of DMN-specific genes.

We further tested SpaCon’s cross-species generalizability by applying it to the marmoset brain, integrating spatial transcriptomics with functional connectivity derived from resting-state functional MRI data ^40^. The model identified spatial domains that bridged anatomically distant but functionally connected regions (**Fig. 6d, S15b**), including one domain substantially overlapped with the default mode-like network (DMN) ^41^. However, our analysis revealed that SpaCon’s domain represented a more precise, transcriptionally-defined network than the canonical DMN. Compared to the DMN, the SpaCon domain exhibited a significantly larger metagene expression difference between the inside and outside of the domain (Cohen’s d=2.99) and was more accurately reconstructed from its metagenes (**Fig. 6e**). Furthermore, when we partitioned the DMN based on its intersections with SpaCon-defined domains, these overlapping subregions showed markedly higher gene-expression correlation with each other than the non-overlapping DMN regions (**Fig. 6f**), demonstrating that SpaCon coherently groups regions by both gene-expression and functional similarity. GO analysis further clarified the functional distinctions between the SpaCon domain and the broader DMN. While shared metagenes were enriched for neuronal projection, reflecting long-range connectivity, SpaCon-specific metagenes were enriched for ion-channel activity, whereas DMN-specific metagenes were enriched for synaptic functions (**Fig. 6g-h**). These results show that SpaCon refines canonical functional networks by defining them on a more precise molecular basis.

In summary, SpaCon is a powerful and versatile framework that effectively integrates gene expression with diverse modes of brain connectivity—from anatomical tracing to calcium imaging and fMRI. Its successful application in both mouse and marmoset demonstrates its robustness and broad cross-species generalizability, providing a scalable tool to uncover biologically meaningful structures in the brain.

## Discussion

By integrating single-cell resolution spatial transcriptomics with whole-brain connectomics, our study establishes a powerful new framework for decoding the principles of brain organization. We first revealed a strong coupling between connection strength and gene-expression profiles in the corticothalamic circuits, especially for glutamatergic neurons (**Fig. 1**). By analyzing the extrasomatic transcriptome, we provided evidence that axonal tracts carry mRNA repertoires that reflect their specific cortical origins, physically bridging the transcriptomes of connected regions (**Fig. 2**). To address the challenge of integrating spatial transcriptomic data with the global connectivity patterns, we developed SpaCon, a scalable and versatile deep-learning method for multi-modal data fusion (**Fig. 3**). SpaCon not only identifies coherent brain domains defined by their dual molecular and connectional identities (**Figs. 4, 6**) but also enhances the classification of neuronal subtypes where transcriptomic data alone is insufficient. (**Fig. 5**). This study provides both a new conceptual understanding of brain organization and a powerful computational tool to explore the complex relationship between the transcriptome and the connectome.

While the association between gene expression and brain connectivity is well-established ^3–10^, our study provides a more refined understanding of this relationship (**Figs. 1, 2**). First, previous studies have not fully distinguished the contributions of different cell types to the transcriptome-connectome relationship in the corticothalamic circuit. Using single-cell resolution spatial transcriptomic data ^27,29^, we demonstrated that cortical-thalamic connectivity patterns primarily correlate with gene expression in thalamic glutamatergic neurons. This aligns well with anatomical evidence that thalamic glutamatergic neurons are the predominant target of cortical projections ^42–45^. Second, our work refines the established principle of spatial autocorrelation, where transcriptional similarity between brain regions typically decays with physical distance ^2,8,14^. We demonstrated that this rule depends on anatomical connection strength. While the distance-decay effect persists for weakly connected regions, strong connectivity maintains high transcriptional similarity between regions, irrespective of physical distance. Finally, our characterization of the corpus callosum provides critical functional validation for the emerging field of spatially resolved extrasomatic mRNAs ^17,18^. Beyond merely confirming their presence or organized distribution, our results demonstrate that these transcripts serve as structured molecular cargoes that encode the identity of their distant somatic origins. This finding not only aligns with established mechanisms of axonal mRNA transport ^19–22^ but also extends current frameworks by showing that these extrasomatic signals constitute a readable physical record of long-range connectivity, offering a novel modality to bridge local molecular profiles with global circuit architecture. These discoveries converge to present a more integrated view, where the brain’s molecular and connectional landscapes are linked through specific cellular and physical pathways.

Systematically linking local molecular data with global circuit architecture requires a new computational approach, as existing algorithms for either multi-omics ^46–49^ or spatial transcriptomics ^23–26,46^ were not tailored for this integrative task. We therefore developed SpaCon, a deep-learning framework with a distinct architecture designed for this challenge (**Fig. 3**). SpaCon employs a dual-encoder graph attention autoencoder that learns features from the local spatial graph and the global connection graph separately before their fusion, allowing it to handle these two heterogeneous data types more effectively than traditional models ^23–26^. To overcome the computational challenge of integrating multi-modal data, SpaCon employs a highly efficient subgraph sampling strategy with an adjustable batch size, ensuring scalability for the analysis of massive, whole-brain datasets. Crucially, a tunable hyperparameter (α) provides analytical flexibility, allowing the model’s final output to emphasize either global connectivity patterns (as α approaches 1) for defining long-range domains (**Figs. 4, 6**) or local spatial context (as α approaches 0) for fine-grained tasks like cell-subtype classification (**Fig. 5**). Consequently, SpaCon not only predicts connectivity from gene expression with greater accuracy but also enhances cell-subtype classification when gene-expression features are ambiguous, establishing it as a powerful and versatile tool for multimodal brain mapping.

While this study establishes a robust framework, its conclusions are drawn primarily from the mouse cortico-thalamic system. A crucial next step is to validate these findings and the SpaCon methodology across diverse neural circuits and developmental stages to establish broader principles. The analysis is also constrained by the technical limitations of current spatial transcriptomics, which cannot unambiguously distinguish axonal mRNA from that of adjacent cells. Future work may address this by using new anterograde tracing methods alongside axon-specific transcriptomic profiling methods, such as Axon-Seq ^50^. Looking ahead, SpaCon’s analytical power can be significantly expanded. Incorporating emerging single-neuron projection data ^51^ would enhance its sensitivity to rare cell types and fine-scale microcircuits, while the inclusion of other data modalities, such as epigenomics, proteomics, and electrophysiology, would yield a more holistic understanding of brain organization.

## Methods

### Spatial transcriptome data

MERFISH dataset. Zhang et al. ^27^ used Multiplexed Error-Robust Fluorescence In Situ Hybridization (MERFISH), a spatially resolved single-cell transcriptomics method, to generate a comprehensive cellular atlas of the adult mouse brain. The dataset comprises expression profiles for more than 1,100 genes in over 9.3 million cells, obtained from coronal and sagittal sections of four adult mice.

Slide-seq dataset. Langlieb et al. ^28^ applied the Slide-seq technique to capture spatial transcriptomic data from contiguous coronal sections of one hemisphere of an adult female mouse brain, with approximately 100 µm spacing between slices. In total, 101 coronal sections were profiled, covering expression of over 35,000 genes in more than 3.7 million cells.

Stereo-seq dataset. Han et al. ^29^ employed Stereo-seq, a sequencing-based whole-genome and high-resolution spatial transcriptomics technology, to collect 123 coronal slices from the mouse brain hemispheres. This dataset contains expression data for approximately 30,000 genes across more than 4 million cells.

### Connectome data

Structural connectivity data. The Allen Mouse Brain Connectivity Atlas ^30^ employs adeno-associated viral vectors expressing enhanced green fluorescent protein (EGFP) to trace axonal projections from specific regions and cell types. High-throughput serial two-photon tomography was subsequently used to image the EGFP-labeled axons throughout the entire brain. By registering each experiment’s results into a common 3D reference space, a comprehensive connectivity matrix of the entire mouse brain was generated.

Mouse widefield calcium imaging data. We utilized the functional connectivity matrix generated by Fei et al. ^36^ from widefield calcium imaging, which had been registered to a 100 µm reference atlas and comprised 7,002 cortical spots. By mapping this matrix to the spatial transcriptomics data, we obtained a functional connectivity matrix for 3,372 spots in the left hemisphere of the mouse brain.

### Data preprocessing

Spatial transcriptomics data. For the MERFISH data ^27^, we used the Zhuang-ABCA-1-raw.h5ad (coronal) and Zhuang-ABCA-3-raw.h5ad (sagittal). For the Slide-seq data ^28^, we used Puck_Num_01.h5ad through Puck_Num_101.h5ad, comprising 101 sections. For the Stereo-seq data ^29^, we used the mouse1 dataset; following the procedures described by Han et al. ^29^, we preprocessed the data and removed genes expressed in fewer than 30 cells and cells expressing fewer than 150 genes. All of the above datasets were normalized and log-transformed, and the top 1500 highly variable genes (HVGs) were selected for downstream analyzes (the MERFISH dataset contains only ~1,100 genes, we used all genes in that case).

Connectome data. The structural connectivity data used in this study were obtained from the Allen Mouse Brain Connectivity Atlas. The connection strength is defined as the ‘normalized projection volume’, which is calculated as the volume of detected projection signals across all voxels within a given structure (in mm^3^), divided by the total volume of detected signals in the manually annotated injection site. The processing pipelines for the Allen Mouse Brain Connectivity data and the mouse widefield calcium imaging data followed the procedures described in Fei et al. ^36^.

Registration of connectome and spatial transcriptome data. We utilized the connectome data from Allen’s neural tracing dataset at a 100-um resolution ^30,31^, while the spatial transcriptomics data ^27–29^ with a higher resolution. To jointly analyze these datasets, we first needed to register the connectome data with the spatial transcriptomics data. Although both were aligned to Allen’s Common Coordinate Framework (CCF) ^52^, their resolutions differed. To address this, we performed a nearest-neighbor search: using the 100-um connectome data as the reference, we matched each cell in the spatial transcriptomics data to its nearest connectome point. As a result, each connectome point corresponded to approximately ten transcriptomic cells. We then aggregated the gene expression data of these cells and calculated the average x and y coordinates. The resulting values provided both gene expression profiles and spatial coordinates for each spot, establishing a one-to-one correspondence between the combined spatial transcriptomics spots and connectome points.

Extracellular gene expression data. Spatial transcriptomics platforms typically employ cell segmentation algorithms to define cell boundaries and attribute the mRNAs within those regions to individual cells. However, mRNAs are also present in the extracellular space. To examine extracellular transcripts specifically, we excluded all intracellular gene expression, retaining only the extracellular component. Using raw MERFISH data from Zhang et al. ^27^, which contains both single-molecule coordinates and segmented cell boundaries, we removed mRNAs falling inside cell boundaries. Then, applying the same nearest-neighbor search, we assigned each extracellular mRNA to its closest cell, labeling it as that cell’s extracellular expression. This procedure allowed direct comparison of intracellular versus extracellular gene expression in our subsequent analyzes.

### Analysis of the relationship between connectivity and gene expression

Clustering of connectivity and gene expression. Based on the mean connection strength from each cortical region to each thalamic nucleus (**Fig. S1c)**, we performed UMAP for dimensionality reduction and clustering of the thalamic nuclei (**Fig. 1a**), yielding eight groups of nuclei. Nuclei within the same group primarily originated from projections of the same cortical area (**Fig. S2a-b**). Next, we applied UMAP to MERFISH spatial transcriptomics data (including both coronal and sagittal datasets) for the thalamic nuclei (**Fig. 1b, S2c-e**). Using published cell-type annotations ^27^, we separately performed dimensionality reduction and clustering for glutamatergic neurons, GABAergic neurons, and non-neuronal cells, and visualized each via UMAP. In the gene-expression UMAP plots, each thalamic nucleus was colored according to its connectivity-defined groups from **Fig. 1a**.

Quantitative analysis of thalamic gene expression (PC1) and corticothalamic mean connectivity. Based on the MERFISH spatial transcriptomics data described above, we performed principal component analysis (PCA) on gene expression profiles of different thalamic cell types with Scanpy ^53^. We extracted 50 principal components and plotted the first two (PC1 and PC2) in PCA scatterplots (**Fig. 1c, S3a-d**). To examine the relationship between each cell’s PC1 score and its cortical input strength, we averaged the corticothalamic connectivity matrix across all cortical spots, thereby obtaining a connectivity vector whose length matches the number of thalamic cells so that each cell is characterized by a PC1 value and an average connection strength value. We then grouped cells by thalamic nucleus, computed mean PC1 and mean connectivity for each nucleus, and calculated Pearson correlation coefficients between PC1 and connection strength for each cell type (**Fig. 1c-d, S3a-d**).

Thalamic gene expression changes with connection strength. We next used scVelo ^32^ to examine how gene expression dynamics vary with connection strength. scVel’s dynamical model fits splicing kinetics to compute a likelihood score for each gene, indicating how strongly its expression dynamics align with connectivity patterns. We ranked genes by this likelihood and selected the top 200 for further analysis (**Fig. 1f**). To quantify expression change, we calculated Euclidean distances between adjacent cells (cell-based trends) or adjacent genes (gene-based trends). To smooth out noise and highlight major trends, we applied a sliding window filter to the distance sequences (window size = 50 for cells; window size = 10 for genes). Using the point of maximal change in the gene-based sequence, we split the 200 genes into weak connectivity associated genes (first 95 genes) and strong connectivity associated genes (last 105 genes), then performed GO enrichment analysis separately on each gene set (**Fig. 1g**).

Correlation analysis of connection strength and gene co-expression. To quantify the relationship between pairwise connection strength and gene-expression correlation for each cortical and thalamic spot, we calculated the gene-expression correlation between cortex and thalamus. However, given that the registered spatial transcriptomics data encompasses approximately 30,000 cortical spots and 5,000 thalamic spots, calculating pairwise correlations would necessitate billions of correlation computations. Furthermore, the prevalence of zero gene counts in many spots could introduce inaccuracies if correlations are calculated directly at the spot level. To mitigate these challenges, we divided each Allen Brain Atlas region into smaller, manageable regions of interest (ROIs). Specifically, we utilized hierarchical clustering based on the spatial coordinates of all spots within each region, ensuring each cluster contained approximately 20 spots, defining these clusters as ROIs. For regions with fewer than 15 spots, clustering was bypassed, and the region was directly designated as a single ROI.

We then conducted analyzes at the ROI level. ROI-to-ROI connectivity was defined as the mean connection strength across all spot pairs between two ROIs, and ROI gene expression was defined as the average expression of all spots within each ROI. We extracted corticothalamic projection data from the Allen neural tracing dataset. Subsequently, we calculated the Pearson correlation between ROI gene-expression profiles for each cortical-thalamic ROI pair. This approach yielded a paired measure of connection strength and gene-expression correlation for every corticothalamic ROI pair. Finally, we visualized their relationship in a scatter plot with connection strength on the x-axis and gene-expression correlation on the y-axis (**Fig. 1h**). To characterize the overall distribution in the scatter plot (**Fig. 1h**, *middle*), we applied kernel density estimation (KDE) to extract its outer contour. Because the raw contour was noisy, we then fitted a fifth-order polynomial to the KDE contour to produce a smoothed boundary (red dashed line in **Fig. 1h**). We calculated the skewness (−0.32) and kurtosis (1.83) of this smoothed contour (a normal distribution has skewness = 0, kurtosis = 3). To formally evaluate normality, we conducted the Jarque-Bera test, yielding JB = 73.96 (p < 0.0005), indicating a significant deviation from a normal distribution.

To address the high density of points and the absence of direct linear correlation between connection strength and gene-expression correlation (i.e., weak connectivity does not necessarily imply low expression correlation), we further quantitatively summarized their relationship. Specifically, we binned the connection strength into 30 equal-width intervals and calculated the mean correlation for all points in each interval (thus representing each bin by a single summary point; **Fig. 1h**, *right*). We replicated this pipeline on the MERFISH (coronal) ^27^, Stereo-seq (coronal) ^29^, and Slide-seq (coronal) ^28^ spatial transcriptomics datasets, obtaining consistent results (**Fig. S3e**).

Regional analysis of layer 5/6 corticothalamic gene expression and connectivity. The connectivity data used above were based on wild-type mice. Previous studies ^31^ provided neural-tracing data at the regional level from three transgenic mouse lines: *Rbp4*-Cre_KL100 (Layer 5), *Ntsr1*-Cre_GN220 (Layer 6), and *Syt6*-Cre_KI148 (Layer 6). Those data describe the strength of projections from cortex to thalamic nuclei. The reference ^31^ reported that projections across these three datasets are correlated, we averaged their connectivity values. We also merged SSs-1 and SSs-2 into SSs, and MOs-1 and MOs-2 into MOs, yielding a 25×44 connectivity matrix (25 cortical regions × 44 thalamic nuclei; **Fig. S4a**). Next, we calculated the gene-expression correlations between the 25 cortical regions and 44 thalamic nuclei (only cortical Layer 5/6 and thalamic glutamatergic neurons were retained). To achieve more accurate correlation calculations, instead of averaging the gene expression across all cells in each region, we applied hierarchical clustering to group cells into ROIs of ~100 cells each, analogous to our earlier approach. For each ROI, we computed the mean gene expression of its constituent cells. We then calculated Pearson correlations between ROI gene-expression profiles for every cortical-thalamic pair. Finally, we averaged these ROI-level correlations to produce a 25 × 44 gene-expression correlation matrix.

Based on the analysis results from wild-type mice, regions with stronger connectivity exhibit more noticeable gene expression similarities. Therefore, we focused on these strongly connected regions and selected unconnected regions for comparison. For each cortical region, we classified thalamic nuclei as ‘connected’ if their log_10_(-normalized projection volume) exceeded −2, and as ‘unconnected’ based on true negative connections identified by Harris et al.^31^. For visualization, we set the log_10_(-normalized projection volume) of true negatives to −10 (**Fig. S4a**). Using the previously generated 25×44 gene-expression correlation matrix, we calculated the average correlation between each cortical region and its ‘connected’ / ‘unconnected’ thalamic nuclei (**Fig. 1i**). Next, to identify genes enriched in both cortex and ‘connected’ nuclei—but depleted in ‘unconnected’ nuclei—we performed differential expression analysis with Scanpy’s *rank_genes_groups* function ^53^. First, we identified genes upregulated in cortex versus ‘unconnected’ nuclei; then we identified genes upregulated in ‘connected’ versus ‘unconnected’ nuclei. The intersection of these two sets yielded our target genes. For each cortical area, we filtered for genes with log_2_ fold change > 2, then calculated the mean expression of these genes in cortex, ‘connected’ nuclei, and ‘unconnected’ nuclei. To standardize across genes, we converted these means to percentages and plotted them in a radar chart (**Fig. S4b**). Finally, we replicated this entire workflow on Stereo-seq dataset (**Fig. S4c–d)**.

### Intracellular and extracellular gene expression analysis of callosal cells

Utilizing the methods described above, we extracted gene-expression data from the extracellular regions of the MERFISH spatial transcriptomic datasets, encompassing both coronal and sagittal sections. Because mRNAs located too far from the cells may introduce errors (e.g., originating from improperly segmented cells), we filtered the extracellular mRNAs for each cell, retaining only the 30% nearest molecules.

We then performed differential expression analysis comparing extracellular versus intracellular transcripts in corpus callosum cells using Scanpy’s *rank_genes_groups* function (**Fig. 2a**). We identified 151 upregulated genes (log_2_ fold change > 1, p < 0.005) and 50 downregulated genes (log_2_ fold change < −1, p < 0.005). Next, we grouped these genes by cell-type enrichment. By merging glutamatergic and GABAergic subtypes into ‘Glut’ and ‘GABA’, respectively, we got six distinct cell types: ‘Glut’, ‘GABA’, ‘Immune’, ‘Vascular’, ‘OPC-Oligo’ (oligodendrocyte precursor cells-oligodendrocytes), and ‘Astro-Epen’ (astrocytes-ependymal). We then calculated the average expression levels of these upregulated and downregulated genes across these six cell types within the cortical corpus callosum region (**Fig. 2a, S6c**).

Additionally, **Fig. S6a** displays each gene’s expression across these six cell types and the results of GO enrichment analysis^54^. Notably, **Fig. 2a** shows that genes downregulated in the extracellular space relative to the intracellular space were highly expressed in GABAergic neurons. Further analysis revealed that several of these genes were co-expressed in both GABAergic neurons and OPC-Oligo cells (**Fig. S6b**).

To show that these results were not affected by random factors, we analyzed the two distinct MERFISH datasets, coronal and sagittal. Among 1,111 genes measured intracellularly and extracellularly in both datasets, we performed differential expression analysis and obtained log_2_ fold change for each gene. We then identified the top 100 genes with the highest log_2_ fold change and the bottom 100 genes with the lowest in each dataset, compared their overlap (78 in the top 100; 83 in the bottom 100; **Fig. 2c**) and calculated the correlation of log_2_ fold change between datasets (**Fig. 2d**). Then we performed GO enrichment analysis on these overlapping genes (**Fig. 2e**).

We also further analyzed the marker genes specific to different layers of the cerebral cortex. We first performed differential expression analysis to identify genes enriched in each cortical layer (**Fig. S8a**). We then calculated the average intracellular and extracellular expression of these marker genes in the callosal region (**Fig. 2f, S8b**).

### Analysis of corticocallosal connectivity and gene expression

To further explore the relationship between cortical projections and extracellular gene expression in the corpus callosum, we focused on MERFISH spatial transcriptomics data from the midline sagittal slice, which captures callosal fibers connecting the cortex to its contralateral counterpart. By averaging the connectivity data from each cortical region to the corpus callosum, we derived the connection strength matrix between each cortical region and every spot within the corpus callosum (**Fig. S8c-d**).

To ensure robust results, we filtered the matrix to retain only regions with substantial axonal projections. Since the corpus callosum data arise exclusively from extracellular axonal transcripts, we removed any row or column in which all connectivity values were < 0.001 (fewer cortical projections). This rigorous filtering process retained only those regions where the cortex exhibited strong connections to the corpus callosum (**Fig. S8e**), specifically 25 cortical regions and three distinct parts of the corpus callosum: the anterior (genu), middle (body), and posterior (splenium). We then clustered the corpus callosum spots based on the filtered connection strength, identifying three clusters that corresponded to the genu, body, and splenium. Finally, we averaged connection strength matrix within each segment to produce a 25 × 3 matrix representing mean connectivity from each cortical region to each callosal part (**Fig. 2g**).

We first divided the cortex into three regions (CTX1, CTX2, CTX3) based on projection patterns to the three callosal areas. Using Scanpy’s *rank_genes_groups* function, we identified genes enriched in each cortical region. We selected the top five genes by log_2_ fold change for each region and calculated their average expression per spot across the three callosal areas (**Fig. 2h, S8f**).

Next, we calculated the gene-expression correlation between the 25 cortical regions and the three extracellular parts of the corpus callosum. As before, we plotted a scatter plot, where the x-axis depicted connection strength and the y-axis represented gene-expression correlation (**Fig. 2i**). Only points with an average connection strength > 0.0001 were included in the plot. We repeated this experiment on the Stereo-seq dataset. However, due to the Stereo-seq dataset only includes coronal sections, we mapped callosal segments along the anteroposterior (AP) axis (z-coordinate). Based on connectome-derived boundaries, we defined z < 43 as genu, z = 50-54.5 as body, and z ≥ 70 as splenium. We then applied the same connectivity-expression workflow to these Stereo-seq segments (**Fig. S8g**).

### The SpaCon framework

SpaCon is an innovative graph neural network framework that integrates connectome and spatial-transcriptome data via graph-attention autoencoders. By fusing these two modalities into a single cohesive graph, SpaCon learns a unified representation. It comprises three main modules (**Fig. 3a**): the Spa-encoder, the Con-encoder, and the decoder. The Spa-encoder ingests the spatial graph, while the Con-encoder processes the connectivity graph. Their output feature embeddings are then concatenated and passed to the decoder, which reconstructs the gene-expression matrix. This reconstruction is used to compute the loss function, allowing SpaCon to be iteratively optimized.

### Constructing the spatial graph and connection graph

We construct the spatial graph and connection graph for the Spa-encoder and Con-encoder respectively, utilizing local spatial information and global connection information.

#### Spatial graph

Following the methodology outlined in STAGATE ^24^, we constructed a 3D spatial graph ***G***_*s*_(***V, E***) using gene-expression and spatial-coordinate data from spatial transcriptomics. Here, ***V*** represents the set of *n* nodes, and ***E*** represents the set of edges. First, on each individual slice, we constructed a 2D spatial neighborhood graph. For a point *S*_*i*_(*x*_0_, *y*_0_) in a slice of the spatial transcriptomics data, its neighbors are all points *S*_*j*_ = (*x, y*) satisfying 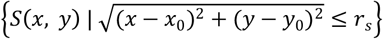, where *r*_*s*_ is a tunable neighborhood radius. Let ***A***_*s*_ ∈ ℝ^*n*×*n*^ be the adjacency matrix constructed from *n* points in the spatial transcriptomics data. If point *S*_*j*_ is located within the neighborhood of point *S*_*i*_ with a radius *r*_*s*_, then ***A***_*s*_ *i, j* = 1; otherwise, ***A***_*s*_ *i, j* = 0. Next, we extended this to adjacent slices: using a radius *r*_*a*_ (chosen to exceed the inter-slice spacing), we connected points across slices. Specifically, if *S*_*i*_ and *S*_*j*_ lie in different slices and *S*_*j*_ − *S*_*i*_ S *r*_*a*_, then ***A***_*s*_ *i, j* = 1; otherwise, ***A***_*s*_ *i, j* = 0.

#### Connection graph

We constructed a 3D connection graph ***G***_*c*_(***V, E***) using connectome data, where ***V*** represents a set of *n* nodes, and ***E*** represents a set of edges. Specifically, by using Allen’s neural tracing data, we obtained a voxel-level (100-um) connection strength matrix. Due to the asymmetry of the connectivity matrix, we symmetrized it by selecting the maximum connection strength between each pair of symmetric positions. Subsequently, we transformed the connection strength matrix into an adjacency matrix, denoted as ***A***_*c*_ ∈ ℝ^*n*×*n*^. To mitigate the impact of noise, we established a threshold of 0.001 for the adjacency matrix. In this matrix, ***A***_*c*_(*i, j*) represents the connection strength between nodes *i* and *j*; if ***A***_*c*_ *i, j* < 0.001, then ***A***_*c*_ *i, j* = 0; if ***A***_*c*_(*i, j*) 0.001, then ***A***_*c*_ *i, j* = 1. After applying the threshold, the sparsity of the resulting adjacency matrix is reduced to approximately 3%. Additionally, we added self-loops to each node, meaning ***A***_*c*_ *i, i* = 1, to obtain the final adjacency matrix ***A***_*c*_.

#### Node features

We use the gene-expression matrix ***E*** ∈ ℝ^*n*×*m*^ from spatial transcriptomics as the node features for both the spatial graph and the connection graph, where *n* is the number of spots and *m* is the number of genes. For the *i*-th node *S*_*i*_, its node feature is the expression level of *m* genes from the *i*-th row of the gene-expression matrix ***E***, denoted as ***E***_{*i*, :}_.

### Graph attention autoencoder

Based on the Graph attention autoencoder proposed by Salehi et al. ^37^, we implemented the construction of the encoder and decoder.

#### Encoder

Both the Spa-encoder and Con-encoder using the same architecture. These encoders transform raw node features into learned representations by aggregating gene-expression vectors from each node’s neighbors. Stacking multiple encoder layers enhances the model’s expressive power. To enable the network to adaptively focus on the most informative neighbors, we integrated an attention mechanism. Following Velickovic et al. ^55^, the attention coefficient between node *i* and its neighboring node *j* in the *k*-th layer of the encoder is as follows:

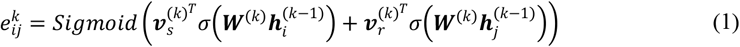

where 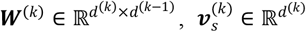 and 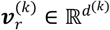 is the trainable parameter of the *k*-th coding layer, *σ* represents the nonlinear activation function, and *Sigmoid* denotes the sigmoid function.

To make the weights of all neighboring nodes comparable, we used the SoftMax function for normalization:

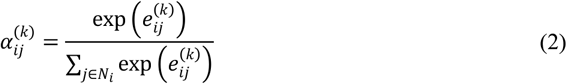

where *N*_*i*_ denotes the set of neighboring nodes of node *i* (including node *i* itself).

The input node features of the encoder are the normalized gene expression matrix. For spot *i*, its node feature is ***x***_*i*_, which is considered as the initial node representation 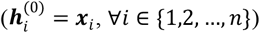. The node representation of node *i* at the *k*-th layer of the encoder is generated by the following formula:

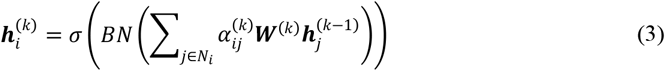

where *BN* denotes Batch Normalization and *σ* represents a non-linear activation function. Assuming the encoder has *L* layers in total, the output of the *L*-th layer is taken as the final node representation:

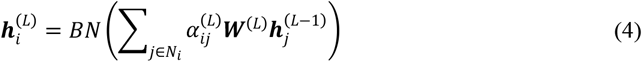

#### Decoder

The decoder mirrors the encoder’s architecture layer by layer but in reverse, receiving the final encoder embeddings and reconstructing node features to enable unsupervised training. In the *k*-th layer of the decoder, the attention coefficient *α* between node *i* and its neighboring node *j* is calculated as follows:

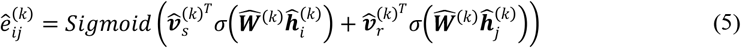

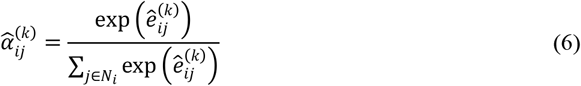

where 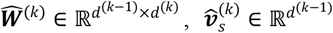, and 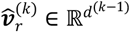 are the trainable parameters in the *k*-th layer of the decoder.

The *k*-th layer of the decoder reconstructs the node feature representation of the (*k* − 1)-th layer as follows:

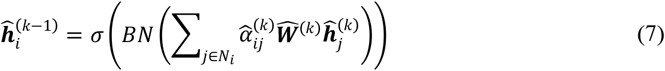

We take the output of the last layer of the decoder as the reconstructed gene expression:

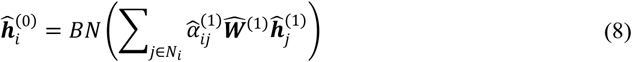

### Training on large-scale datasets through neighbor sampling

Because the connectivity data are information-rich and the transcriptomics data have high resolution, the combined graph becomes extremely large. To address the difficulty of training on such a giant graph, we applied a neighbor-based subgraph sampling method for mini-batch processing so that the model can be trained on GPU. The core idea is to start from a set of target nodes and recursively sample their neighbor layer by layer to assemble a subgraph. In our experiments, we sampled neighbors across three layers: for each target node we first sample 20 first-hop neighbors, then sample 10 second-hop neighbors from those, and finally 10 third-hop neighbors. For comparison, we also implemented random subgraph sampling, where all spots are randomly and evenly partitioned into 64 batches, and each batch’s nodes form a subgraph.

### Co-training of Spa-encoder and Con-encoder

We constructed the spatial graph from local spatial information and the connection graph from global connectivity information. In conventional autoencoders, each graph would require separate model training, which is time-consuming. Additionally, training the two graphs independently makes feature fusion challenging. To overcome these challenges, we employed a co-training strategy, allowing simultaneous training of both encoders and on-the-fly feature fusion. Specifically, we sampled subgraphs only from the connection graph. Each mini-batch is fed into the Con-encoder for training, while the Spa-encoder retrieves a subgraph from the spatial graph using the same set of nodes. Thus, for each batch, both encoders process identical nodes but different edge sets—spatial edges for the Spa-encoder and connectivity edges for the Con-encoder.

During model training, to fuse the node embeddings generated by the two encoders, we either concatenated or element-wise summed the outputs of the Spa-encoder (denoted as ***h***_*s*_ ∈ ℝ^*n*×*p*^) and the Con-encoder (denoted as ***h***_*c*_ ∈ ℝ^*n*×*q*^) to obtain the final output (denoted as ***h*** ∈ ℝ^*n*× (*p*+*q*)^):

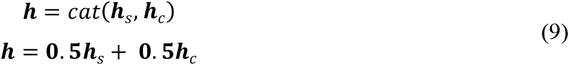

where *cat*(·) denotes feature concatenation. The resulting ***h*** is then passed to the decoder. When we use feature concatenation, the dimension of ***h*** equals the sum of the dimensions of ***h***_*s*_ and ***h***_*c*_, the structure of the decoder is not fully symmetrical to the encoder. Specifically, the decoder’s first layer input size matches the combined dimension of the Spa-encoder and Con-encoder final layers.

### Loss function

Our objective is to learn meaningful node embeddings in an unsupervised manner. We employ an autoencoder architecture to reconstruct node features, thereby encouraging the encoder to capture informative representations. The loss function is defined as follows:

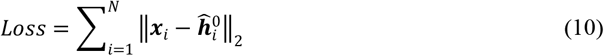

### Clustering

We derive the low-dimensional embeddings from the trained Spa-encoder and Con-encoder to identify spatial domains. After training, the Spa-encoder and Con-encoder produce two embedding matrices, which, after normalization, are ***H***_*s*_ and ***H***_*c*_, respectively. We set a parameter *α* to balance their contributions:

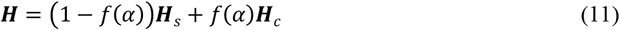

where 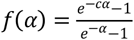, here *c* is a constant, which we set to 4. Here *f* (*α*) is a nonlinear weighting function that makes the clustering results transition more smoothly when *α* values are between 0 and 1. The combined embedding ***H*** is then used to identify spatial domains.

### Implementation details of SpaCon

For the whole-brain spatial transcriptomics dataset, we select the top 1,500 highly variable genes (no gene filtering was needed for MERFISH, which has 1,122 genes). We then performed logarithmic transformation and normalization on the gene expression data. To enhance the model’s fitting capability, both the Spa-encoder and Con-encoder employ a three-layer graph attention network with layer dimensions [*m*, 256, 64, 16], where *m* is the number of input genes. If the features obtained from the two encoders are concatenated, the decoder’s layer dimensions are [32, 64, 256, *m*]. During training, we sampled three hops of neighbors per batch with sizes [20, 10, 10], and used a batch size of 32. We optimized the reconstruction loss with the Adam optimizer (initial learning rate 1×10^−4^, weight decay 1×10^−5^), and employed the Exponential Linear Unit (ELU) activation.

### Analysis of cortex and thalamus clustering results

Clustering and connectivity analysis. To quantify the relationship between clustering results and connectivity, we conducted a statistical analysis. Taking the clustering results between the thalamus and cortex as an example, we calculated the clustering relationships between various cortical regions and thalamic nuclei (**Fig. S12a**). Specifically, for each cortical region, we identified its primary spatial domain—the one containing > 60% of that region’s spots. We then calculated the proportion of spots from each thalamic nucleus that fell within this domain relative to that nucleus’s total spots. This metric captures the association between each cortical region and thalamic nucleus in the clustering output.

Clustering and gene expression analysis. To illustrate the relationship between clustering results and gene expression, we again focus on cortex-thalamus pairs. For each cortical spot *i*, we extracted its connected thalamic neighbors ***V*** (connection strength > 0.001). Within ***V***, we partitioned spots into ***V***_*p*_ (those in the same spatial domain as spot *i*) and ***V***_*n*_ (those in different domains). Both ***V***_*p*_ and ***V***_*n*_ are connected to spot *i*, but only ***V***_*p*_ belongs to the same spatial domain as spot *i*. To explore the relationship between their gene expressions, we calculated the Pearson correlation of gene-expression between spot *i* and each spot in ***V***_*p*_ and ***V***_*n*_, respectively (**Fig. 4b**, *left*). We then conducted a statistical analysis for all points in the cortex. Specifically, we analyzed the gene-expression correlation between each cortical spot *i* and the spots in its corresponding sets ***V***_*p*_ and ***V***_*n*_ (**Fig. 4b**, *right*).

Gene expression predicts connectivity. SpaCon’s clustering partitioned the cortex and thalamus into several spatial domains, most of which contained both cortical and thalamic regions. We selected the spatial domains that contained both cortex and thalamus and then split them into cortical and thalamic subsets. There was a high correlation between thalamic gene-expression PC1 and corticothalamic connection strength (**Fig. 1c**). To investigate whether this relationship is preserved within each SpaCon-derived domain, we built independent linear regression models for every cortical-thalamic subset pair, using the thalamic domain’s PC1 to predict connection strengths from all cortical subsets. We evaluated the model’s prediction error via cross-validation (five-fold if sample size ≥ 5, otherwise leave-one-out), using median absolute percentage error (MDAPE). We then transformed the error matrix into a standardized prediction accuracy matrix by taking the reciprocal (1 / error) and row-normalizing (**Fig. 4c, S13a**). This matrix visually reveals each thalamic domain’s relative predictive ability across all cortical domains.

PLS for identifying metagene in each spatial domain. To identify key genes associated with specific spatial domains, we applied Partial Least Squares (PLS) regression ^38^. We extracted the gene-expression matrix ***X*** and the SpaCon-derived domain labels ***y***. To focus on one target domain, we binarized ***y*** by assigning 1 to the domain of interest and 0 to all others. After training the PLS model, we ranked genes by the absolute value of their PLS coefficients ***γ***, which quantify each gene’s importance in distinguishing the target domain. We then selected the top *k* genes to form a metagene using their expression matrix ***X***_*k*_. The expression of the metagene can be represented as:

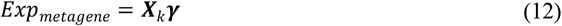

### Cell type classification

Based on spatial transcriptomics data annotated by Zhang et al. ^27^, we benchmarked several clustering algorithms. We selected three layer 6 corticothalamic glutamatergic subtypes, originally labeled ‘0114 L6 CT CTX Glut_1’, ‘0115 L6 CT CTX Glut_2’, and ‘0118 L6 CT CTX Glut_5’. We then renamed these as ‘L6 CT CTX VISp Glut’, ‘L6 CT CTX MO Glut’, and ‘L6 CT CTX ORB Glut’ according to their anatomical locations. These represent the principal corticothalamic glutamatergic neuron subtypes in cortical layer 6 (**Fig. 5a**). To integrate connectivity data, we extracted cells of these three subtypes together with all thalamic cells as our benchmark dataset. We then down-sampled the spatial transcriptomics data to match the resolution of the connectivity data. For SpaCon, clustering was performed on data from all slices. Because most other algorithms do not support 3D clustering, we applied them to a single representative slice and compared their results to SpaCon’s output (**Fig. 5b-d**). We used each method’s recommended parameter settings and ran SpaGCN without histological images.

### Multi-species and multi-modal data analysis

Joint analysis of mouse spatial transcriptomics data and large-scale functional connectivity maps derived from wide-field calcium imaging. We projected the whole-brain coronal spatial transcriptomics data^27^ onto a dorsal-view atlas to align it with wide-field calcium imaging data. The calcium imaging data were processed as described in earlier work ^36^ to yield a functional connectivity matrix. Because these connections were relatively weak and long-range functional connections were not prominent, we applied several preprocessing steps. First, we weighted the original functional connectivity matrix by physical distance: we computed the Euclidean distance between all pairs of nodes and adjusted each connection strength by an exponential function (*weight* = *e*^0.006×*distance*^), so that more distant connections received higher weights. Next, to sparsify the matrix and retain only core connections, we employed a two-stage filtering strategy: we removed all connections with strength < 0.8, then for each node we kept only the top 10 percent of its remaining connections by strength. The resulting sparse matrix had approximately 8.2 percent nonzero entries. We then constructed the spatial graph and the connectivity graph from the processed data for model training.

Joint analysis of marmoset spatial transcriptomics and resting-state fMRI data. We registered both the marmoset spatial transcriptomics and resting-state fMRI data to the marmoset cortical surface ^56^, producing co-registered gene-expression and functional-connectivity maps. For gene expression, we selected the top 3,000 highly variable genes. To build the spatial graph, we connected each point to its 10 nearest neighbors in Euclidean space. For the connectivity graph, we zeroed out weak functional connections and retained only the stronger connections, resulting in a matrix with approximately 9 percent nonzero entries. We then trained SpaCon with α = 1 on these graphs, yielding the final cortical clustering.

## Data availability

The raw data used in this study can be accessed as follows: MERFISH dataset of mouse brain is available at https://alleninstitute.github.io/abc_atlas_access/descriptions/Zhuang_dataset.html ^27^. Slide-seq dataset of mouse brain is available at https://www.braincelldata.org/ ^28^. Stereo-seq dataset of mouse brain is available at https://doi.org/10.12412/BSDC.1699433096.20001 ^29^. Allen structural connectivity data is available at https://connectivity.brain-map.org/ ^30^. MERFISH cell segmentation and decoded results are available at https://download.brainimagelibrary.org/29/3c/293cc39ceea87f6d/processed_data/ ^27^. The wide-field imaging data is available at https://doi.org/10.57760/sciencedb.17064 ^36^. The processed data is available at https://spacon-results-reproduction.readthedocs.io/en/latest/.

## Code availability

SpaCon is implemented in Python and can be accessed through GitHub: https://github.com/quhaichao/SpaCon. The step-by-step tutorial of SpaCon is available at https://spacon-tutorials.readthedocs.io/en/latest/. The source code necessary to reproduce all experiments is available at https://spacon-results-reproduction.readthedocs.io/en/latest/.

## Acknowledgments

The study was supported by grants from the National Science and Technology Innovation 2030 Major Program (Grant No. STI2030-2022ZD0205000 to C.L.), the National Natural Science Foundation of China (32171088 and 32427802 to C.L., U23A20335 and 42271315 to S.Z., 61936007 and 62136004 to J.H.), Shanghai Key Laboratory of Child Brain and Development (No. 24dz2260100 to C.L.).

## Author contributions

C.L. and S.Z. designed the study; H.Q. and W.L. developed the methods; H.Q., W.L., and Y.F. analyzed the data; H.Q., C.L., and W.L. wrote the manuscript; C.L. and S.Z. edited the manuscript. C.L., S.Z. and J.H. supervised the study.

## Competing interests

Authors declare no competing interests.

## Notes

### Competing Interest Statement

The authors have declared no competing interest.

### Summary of Updates

The Methods section has been further improved, and some of the latest references have been added.

