## Supplementary Figures for "Integrative Analysis of Spatial Transcriptome and Connectome by SpaCon"

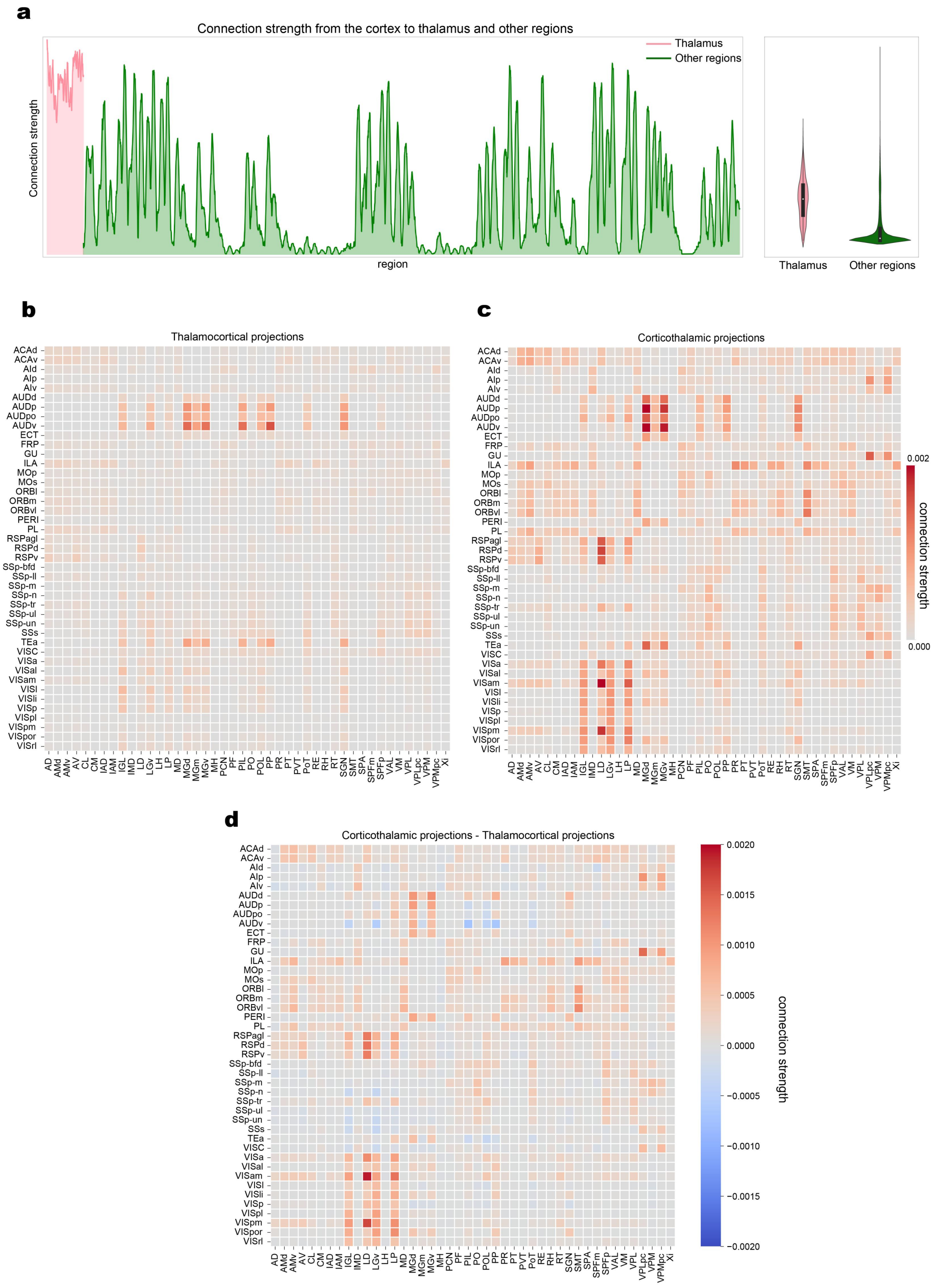


**Supplementary Figure S1: Corticothalamic connections show the strongest projections among all cortical connections.**

**(a)** The strength of projection from the cortex to various brain regions. The left panel shows the connection strengths from the cortex to other brain regions, excluding the cortex itself, based on neural tracing data from wild-type mice of Allen Mouse Brain Connectivity Atlas [1]. The pink-shaded area represents connections to the thalamus, while the green-shaded area represents connections to other brain regions. The right panel shows a violin plot illustrating the distribution of connection strengths from the cortex to the thalamus (pink) and to other brain regions (green).

**(b)** The heatmap shows the connection strength from each thalamic nucleus to each cortical region, with strength values obtained by averaging across each region.

**(c)** The heatmap shows the connection strength from each cortical region to each thalamic nucleus, with strength values obtained by averaging across each region.

**(d)** The differential heatmap shows the difference between the matrix shown in panel (c) and the matrix in panel (b). Positive values are indicated in red, and negative values are indicated in blue.

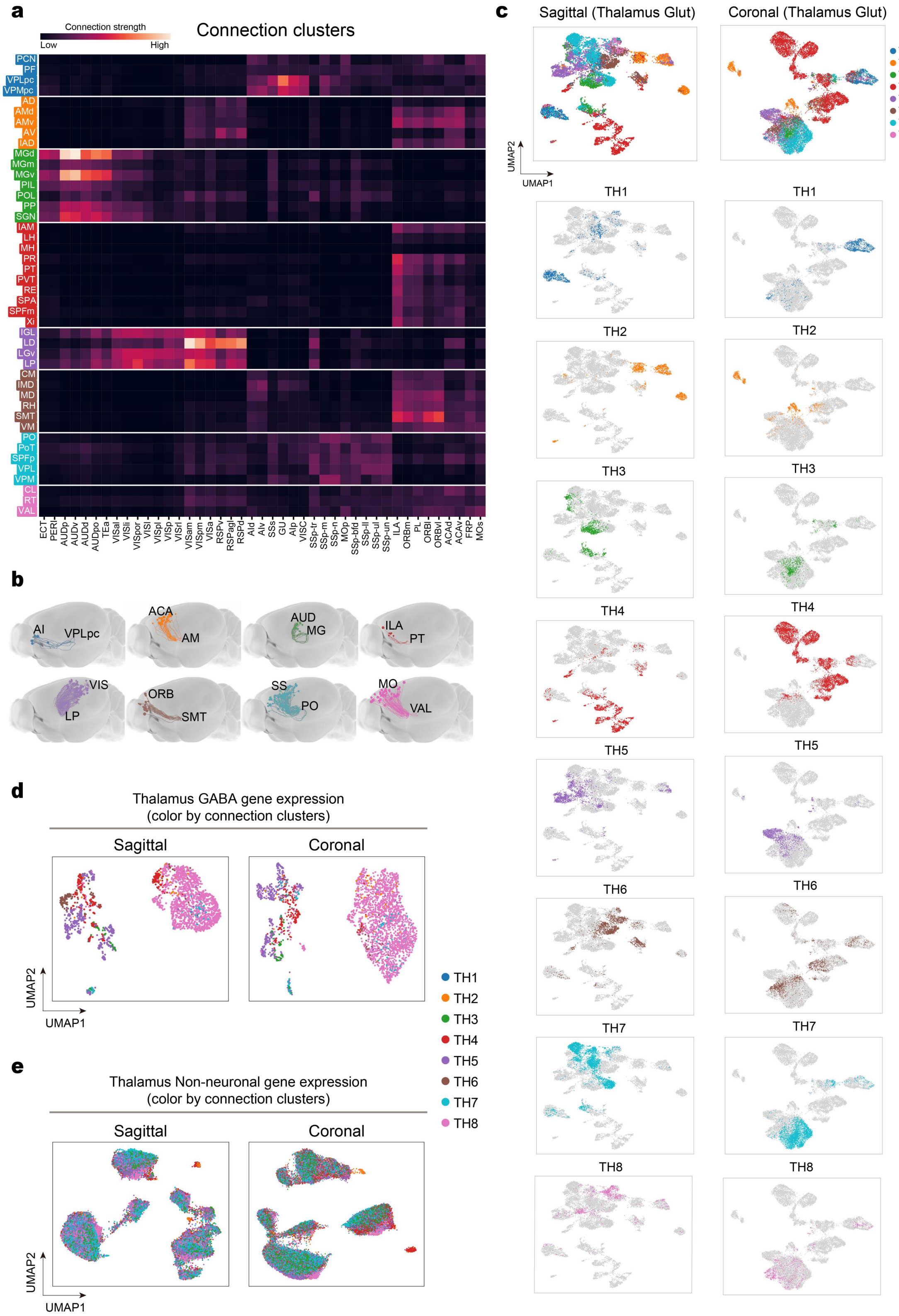

a
Connection strength
Low
High
Connection clusters
PCN PF VPLpc VPMpc AD AMd AMv AV IAD MGd MGm MGv PIL POL PP SGN IAM LH MH PR PT PVT RE SPA SPFm Xi IGL LD LGv LP CM IMD MD RH SMT VM PO PoT SPFp VPL VPM CL RT VAL
ECT PERI AUDp AUDv AUDd AUDpo TEa VISal VISl VISpor VISli VISpl VISp VISrl VISam VISpm VISa RSPv RSPagl RSPd AId AIv SSs GU AIp VISC SSp-tr SSp-m SSp-n MOp SSp-bfd SSp-ll SSp-ul SSp-un ILA ORBm PL ORBl ORBvl ACAd ACAv FRP MOs
b
AI
VPLpc
ACA
AM
AUD
MG
ILA
PT
VIS
LP
ORB
SMT
SS
PO
MO
VAL
c
Sagittal (Thalamus Glut)
Coronal (Thalamus Glut)
TH1 TH2 TH3 TH4 TH5 TH6 TH7 TH8
UMAP2
UMAP1
TH1
TH2
TH3
TH4
TH5
TH6
TH7
TH8
d
Thalamus GABA gene expression
(color by connection clusters)
Sagittal
Coronal
UMAP2
UMAP1
TH1 TH2 TH3 TH4 TH5 TH6 TH7 TH8
e
Thalamus Non-neuronal gene expression
(color by connection clusters)
Sagittal
Coronal
UMAP2
UMAP1


19

**Supplementary Figure S2: Clustering of cortex-to-thalamus connections and UMAP of thalamic gene expression.**

**(a)** Clustering results of the corticothalamic connectivity matrix. The heatmap is organized according to the eight thalamic categories identified in **Fig. 1a**. The matrix represents the connection strength from cortex to thalamus.

**(b)** Examples of projections for each of the eight clusters, with dots representing virus injection sites from Allen Brain Atlas [1].

**(c)** The UMAP clustering of the glutamatergic neurons of thalamic nuclei based on the gene expression features of MERFISH data [2]. The left panels display results from sagittal sections, while the right panels show results from coronal sections. The first row shows all thalamic glutamatergic neurons colored according to the connectivity-based categories shown in **Fig. 1a**, while the bottom panels separately highlight each of the eight connectivity-based categories (colored dots represent the neurons in the respective category, and gray dots represent neurons in remaining nuclei).

**(d)** The UMAP clustering of the GABAergic neurons of thalamic nuclei based on the gene expression features of sagittal and coronal MERFISH data [2].

**(e)** The UMAP clustering of the non-neuronal cells of thalamic nuclei based on the gene expression features of sagittal and coronal MERFISH data [2].

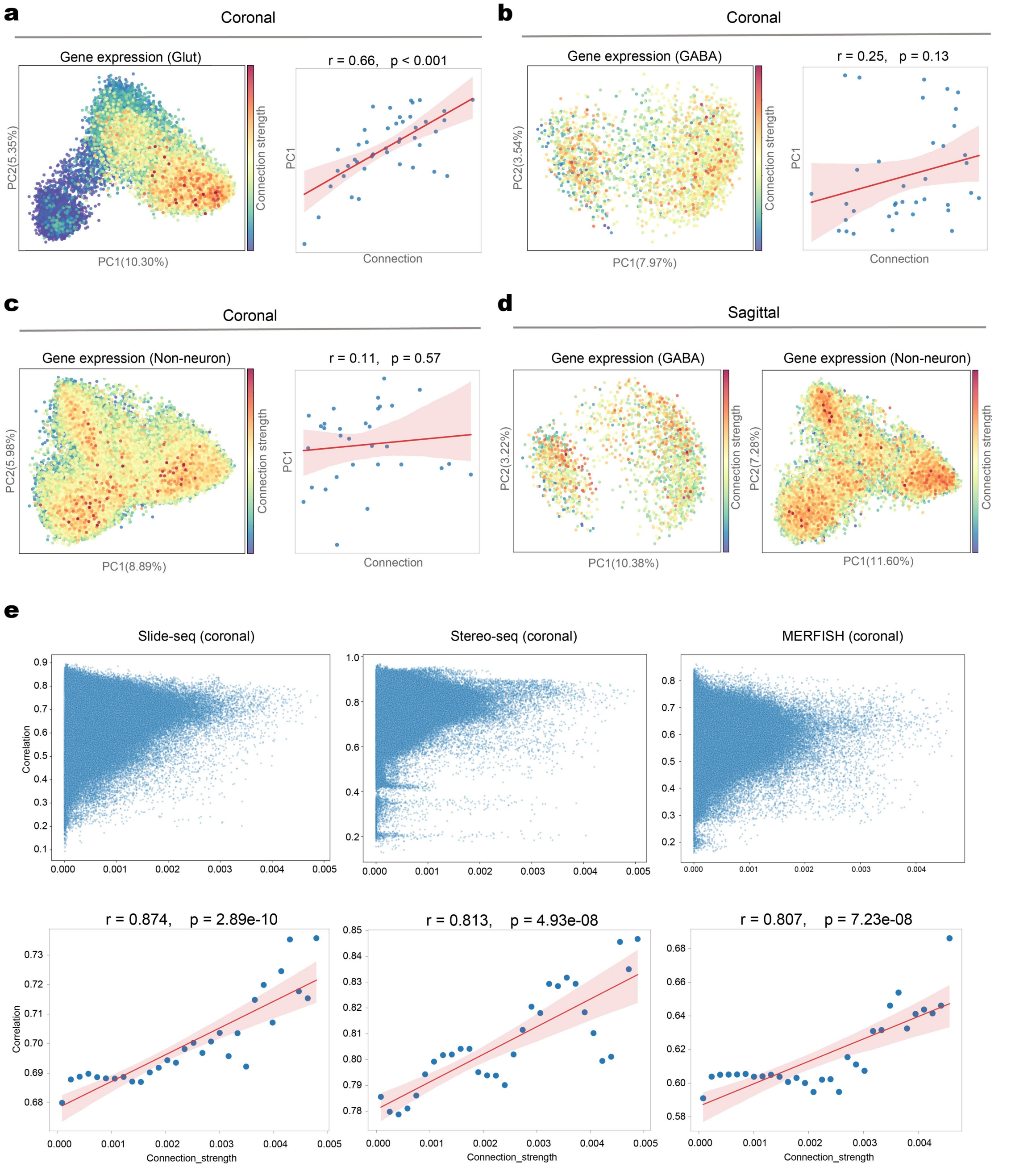


**Supplementary Figure S3: Relationship between gene expression correlation and connection strength of corticothalamic connections.**

**(a)** Relationship between cortical-to-thalamic connection strength and PC1 of gene expression in thalamic glutamatergic neurons (coronal MERFISH data). In the left panel, each dot in the PCA plot represents a single thalamic glutamatergic neuron, colored according to its mean cortical-to-thalamic connection strength. In the right panel, the scatter plot shows the correlation between PC1 and connection strength. Each point corresponds to a thalamic nucleus; the red line is the linear regression fit across all nuclei, and the shaded region indicates the 95% confidence interval. *r* denotes the Pearson correlation coefficient, and *p* the associated P-value.

**(b)** Similar to (a), but using only thalamic GABAergic neurons.

**(c)** Similar to (a), but using only thalamic non-neuronal cells.

**(d)** PCA scatter plots corresponding to **Fig. 1d** and **Fig. 1e**, based on sagittal MERFISH data [2].

**(e)** The relationship between corticothalamic gene expression correlation and connection strength based on Stereo-seq [3], Slide-seq [4], and MERFISH [2], respectively. Transcriptomic data were downsampled to 100 µm resolution to match the connectivity data (**see Methods)**. To reduce computational complexity, 20 spots were further averaged for the correlation analysis. In the top scatter plots, each point represents the connection strength (horizontal axis) between 20 spots in the cortex and 20 spots in the thalamus, and the Pearson correlation of their gene expression patterns (vertical axis). The bottom scatter plots provide a statistical analysis of the top scatter plots by dividing the connection strength into 30 equal intervals. Data from all spots within each interval are averaged to form one point in the bottom plot. The red line is the fitted regression line, with the shaded area indicating the 95% confidence interval.

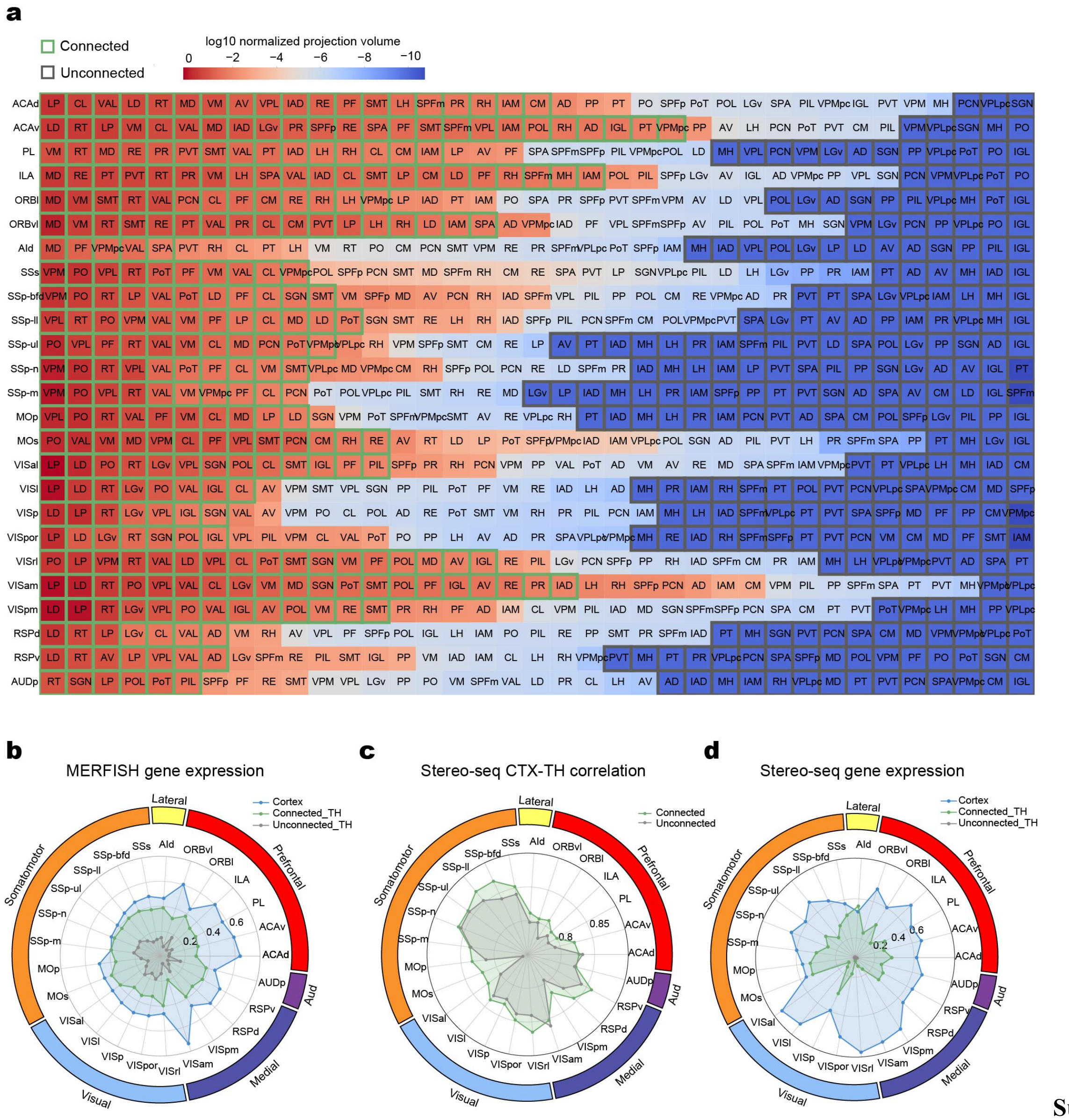


**Supplementary Figure S4: Gene expression correlations between cortical areas and thalamic nuclei (connected and unconnected).**

**(a)** Supplementary information on connected and unconnected thalamic nuclei. This panel shows the specific connected and unconnected thalamic nuclei corresponding to each cortical area. The green boxes highlight thalamic nuclei that are connected to the corresponding cortical region ($\log_{10}$ normalized projection volume > -2), while the gray boxes highlight unconnected nuclei ($\log_{10}$ normalized projection volume < -10). Each cell's color represents the average connection strength between cortical regions and thalamic nuclei in three transgenic mouse lines: *Rbp4*-Cre_KL100 (layer-5 specific), *Ntsr1*-Cre_GN220 (layer-6 specific), and *Syt6*-Cre_KI148 (layer-6 specific) [5], with each row ordered from highest to lowest connection strength.

**(b)** DEGs with higher expression in cortical areas and connected thalamic nuclei vs. unconnected nuclei. The radar chart shows the average expression of these DEGs in the three regions corresponding to 'Cortex' (blue), 'Connected_TH' (green), and 'Unconnected_TH' (gray).

**(c)** Gene-expression correlation between cortical regions and connected (green) vs. unconnected (gray) thalamic areas. Similar to **Fig. 1i**, but based on the Stereo-seq dataset [3].

**(d)** Similar to (b), but based on the Stereo-seq dataset [3].

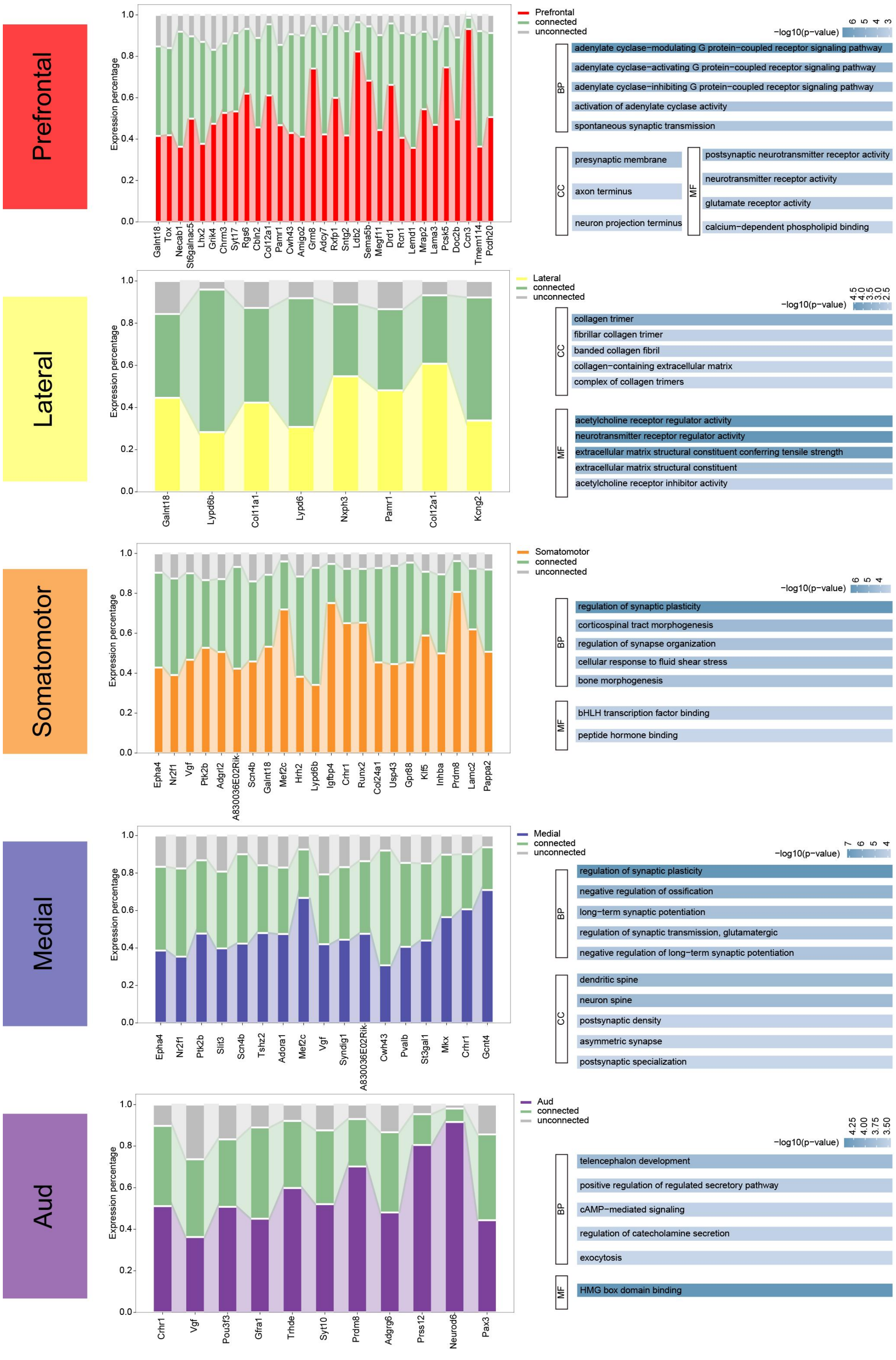

Prefrontal
Expression percentage
1.0
0.8
0.6
0.4
0.2
0.0
Galnt18 Tox Necab1 St6galnac5 Lhx2 Grik4 Chrm3 Syt17 Rgs6 Cbln2 Col12a1 Pamr1 Cwh43 Amigo2 Grm8 Adcy7 Rxfp1 Sntg2 Ldb2 Sema5b Megf11 Drd1 Rcn1 Lemd1 Mrap2 Lama3 Pcsk5 Doc2b Ccn3 Tmem114 Pcdh20
Prefrontal
connected
unconnected
−log10(p−value)
6 5 4 3
BP
adenylate cyclase−modulating G protein−coupled receptor signaling pathway
adenylate cyclase−activating G protein−coupled receptor signaling pathway
adenylate cyclase−inhibiting G protein−coupled receptor signaling pathway
activation of adenylate cyclase activity
spontaneous synaptic transmission
CC
presynaptic membrane
axon terminus
neuron projection terminus
MF
postsynaptic neurotransmitter receptor activity
neurotransmitter receptor activity
glutamate receptor activity
calcium−dependent phospholipid binding
Lateral
Galnt18 Lypd6b Col11a1 Lypd6 Nxph3 Pamr1 Col12a1 Kcng2
Lateral
connected
unconnected
−log10(p−value)
4.5 4.0 3.5 3.0 2.5
CC
collagen trimer
fibrillar collagen trimer
banded collagen fibril
collagen−containing extracellular matrix
complex of collagen trimers
MF
acetylcholine receptor regulator activity
neurotransmitter receptor regulator activity
extracellular matrix structural constituent conferring tensile strength
extracellular matrix structural constituent
acetylcholine receptor inhibitor activity
Somatomotor
Epha4 Nr2f1 Vgf Ptk2b Adgrl2 A830036E02Rik Scn4b Galnt18 Mef2c Hrh2 Lypd6b Igfbp4 Crhr1 Runx2 Col24a1 Usp43 Gpr88 Klf5 Inhba Prdm8 Lamc2 Pappa2
Somatomotor
connected
unconnected
−log10(p−value)
6 5 4
BP
regulation of synaptic plasticity
corticospinal tract morphogenesis
regulation of synapse organization
cellular response to fluid shear stress
bone morphogenesis
MF
bHLH transcription factor binding
peptide hormone binding
Medial
Epha4 Nr2f1 Ptk2b Slit3 Scn4b Tshz2 Adora1 Mef2c Vgf Syndig1 A830036E02Rik Cwh43 Pvalb St3gal1 Mkx Crhr1 Gcnt4
Medial
connected
unconnected
−log10(p−value)
7 6 5 4
BP
regulation of synaptic plasticity
negative regulation of ossification
long−term synaptic potentiation
regulation of synaptic transmission, glutamatergic
negative regulation of long−term synaptic potentiation
CC
dendritic spine
neuron spine
postsynaptic density
asymmetric synapse
postsynaptic specialization
Aud
Crhr1 Vgf Pou3f3 Gfra1 Trhde Syt10 Prdm8 Adgrg6 Prss12 Neurod6 Pax3
Aud
connected
unconnected
−log10(p−value)
4.25 4.00 3.75 3.50
BP
telencephalon development
positive regulation of regulated secretory pathway
cAMP−mediated signaling
regulation of catecholamine secretion
exocytosis
MF
HMG box domain binding


82

**Supplementary Figure S5. Additional results for enrichment analysis of differentially expressed genes (DEG) across cortical modules.**

Stacked bar plots show the percentage expression of DEGs that are upregulated in cortical modules and connected thalamic regions (green) relative to unconnected thalamic regions (gray) across five cortical modules, excluding the visual region that is shown in **Fig. 1i**. The right panels display GO enrichment analysis results for each set of DEGs, categorized into Biological Process (BP), Cellular Component (CC), and Molecular Function (MF). The intensity of shading represents the p-value, with darker shades indicating higher significance.

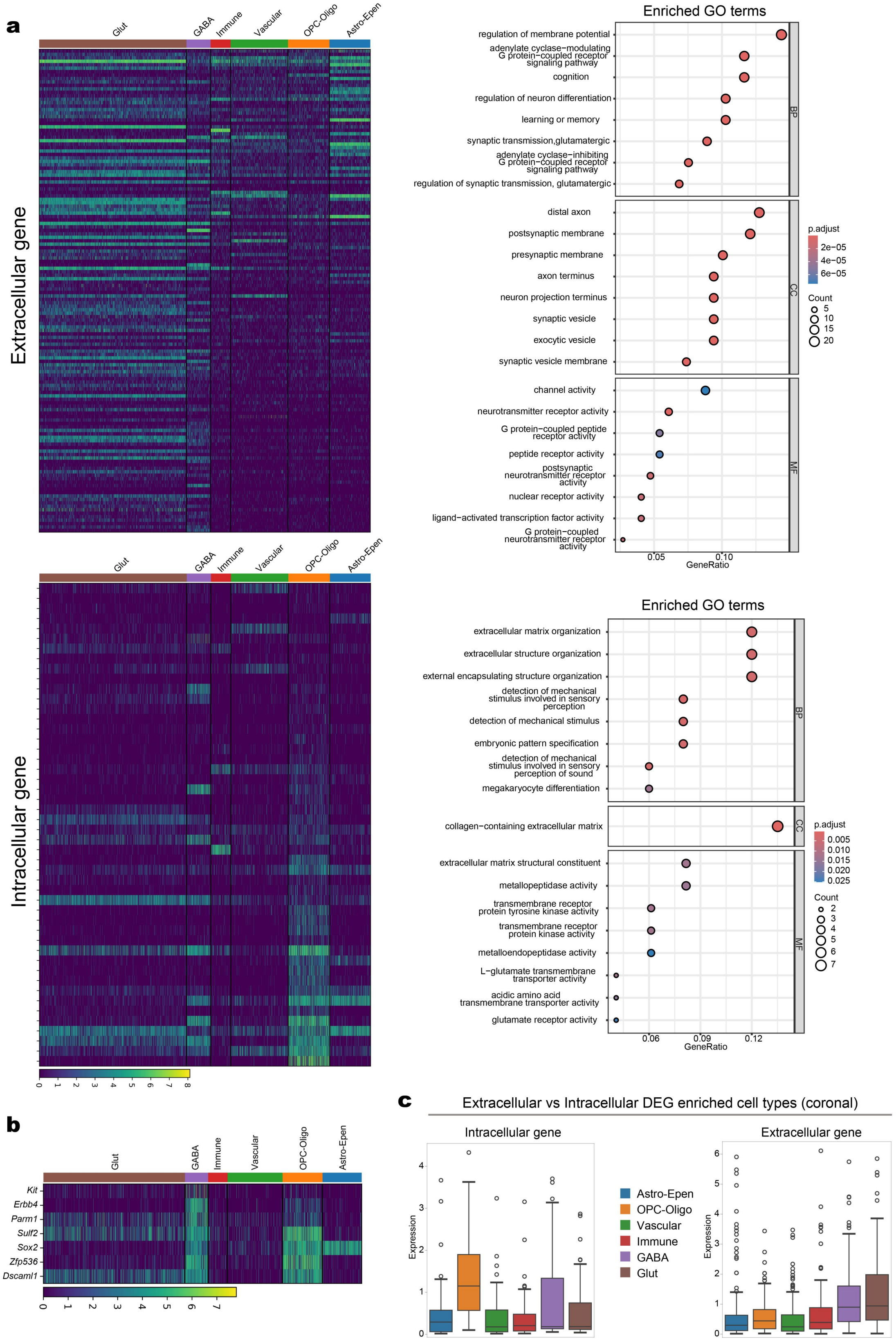

a
Glut
GABA
Immune
Vascular
OPC-Oligo
Astro-Epen
Extracellular gene
Intracellular gene
Enriched GO terms
regulation of membrane potential
adenylate cyclase−modulating G protein−coupled receptor signaling pathway
cognition
regulation of neuron differentiation
learning or memory
synaptic transmission,glutamatergic
adenylate cyclase−inhibiting G protein−coupled receptor signaling pathway
regulation of synaptic transmission, glutamatergic
distal axon
postsynaptic membrane
presynaptic membrane
axon terminus
neuron projection terminus
synaptic vesicle
exocytic vesicle
synaptic vesicle membrane
channel activity
neurotransmitter receptor activity
G protein−coupled peptide receptor activity
peptide receptor activity
postsynaptic neurotransmitter receptor activity
nuclear receptor activity
ligand−activated transcription factor activity
G protein−coupled neurotransmitter receptor activity
BP
CC
MF
p.adjust
2e−05
4e−05
6e−05
Count
5
10
15
20
0.05
0.10
GeneRatio
Enriched GO terms
extracellular matrix organization
extracellular structure organization
external encapsulating structure organization
detection of mechanical stimulus involved in sensory perception
detection of mechanical stimulus
embryonic pattern specification
detection of mechanical stimulus involved in sensory perception of sound
megakaryocyte differentiation
collagen−containing extracellular matrix
extracellular matrix structural constituent
metallopeptidase activity
transmembrane receptor protein tyrosine kinase activity
transmembrane receptor protein kinase activity
metalloendopeptidase activity
L−glutamate transmembrane transporter activity
acidic amino acid transmembrane transporter activity
glutamate receptor activity
p.adjust
0.005
0.010
0.015
0.020
0.025
Count
2
3
4
5
6
7
0.06
0.09
0.12
GeneRatio
b
Kit
Erbb4
Parm1
Sulf2
Sox2
Zfp536
Dscaml1
c
Extracellular vs Intracellular DEG enriched cell types (coronal)
Intracellular gene
Extracellular gene
Expression
Astro-Epen
OPC-Oligo
Vascular
Immune
GABA
Glut


93

**Supplementary Figure S6: Intracellular and extracellular differential gene expression and GO enrichment analysis in the corpus callosum.**

**(a)** Expression levels and GO enrichment analysis of DEGs between intracellular and extracellular compartments in the corpus callosum (**Fig. 2a**). The top heatmap and bubble plot show genes upregulated extracellularly relative to intracellularly, while the bottom show genes downregulated. The heatmaps show the expression levels of all DEGs across various cell types in the cortex, with each row representing a gene and columns representing six cell types: glutamatergic neurons (brown), GABAergic neurons (purple), immune cells (red), vascular cells (green), oligodendrocyte precursor cells-oligodendrocytes (orange), and astrocytes-ependymal cells (blue). The bubble plots display the GO enrichment analysis of the corresponding genes, with bubble color indicating the adjusted p-value, bubble size representing the number of genes enriched in each pathway. The x-axis representing the proportion of enriched genes among the DEGs.

**(b)** Co-expression of genes in GABAergic neurons and oligodendrocyte precursor cells-oligodendrocytes. This heatmap highlights selected genes that are downregulated extracellularly relative to intracellularly in the corpus callosum and are primarily expressed in GABAergic neurons and oligodendrocyte precursor cells-oligodendrocytes.

**(c)** Differential expression analysis of intracellular and extracellular compartments of the corpus callosum. Similar to **Fig. 2a**, but based on the coronal MERFISH dataset [2].

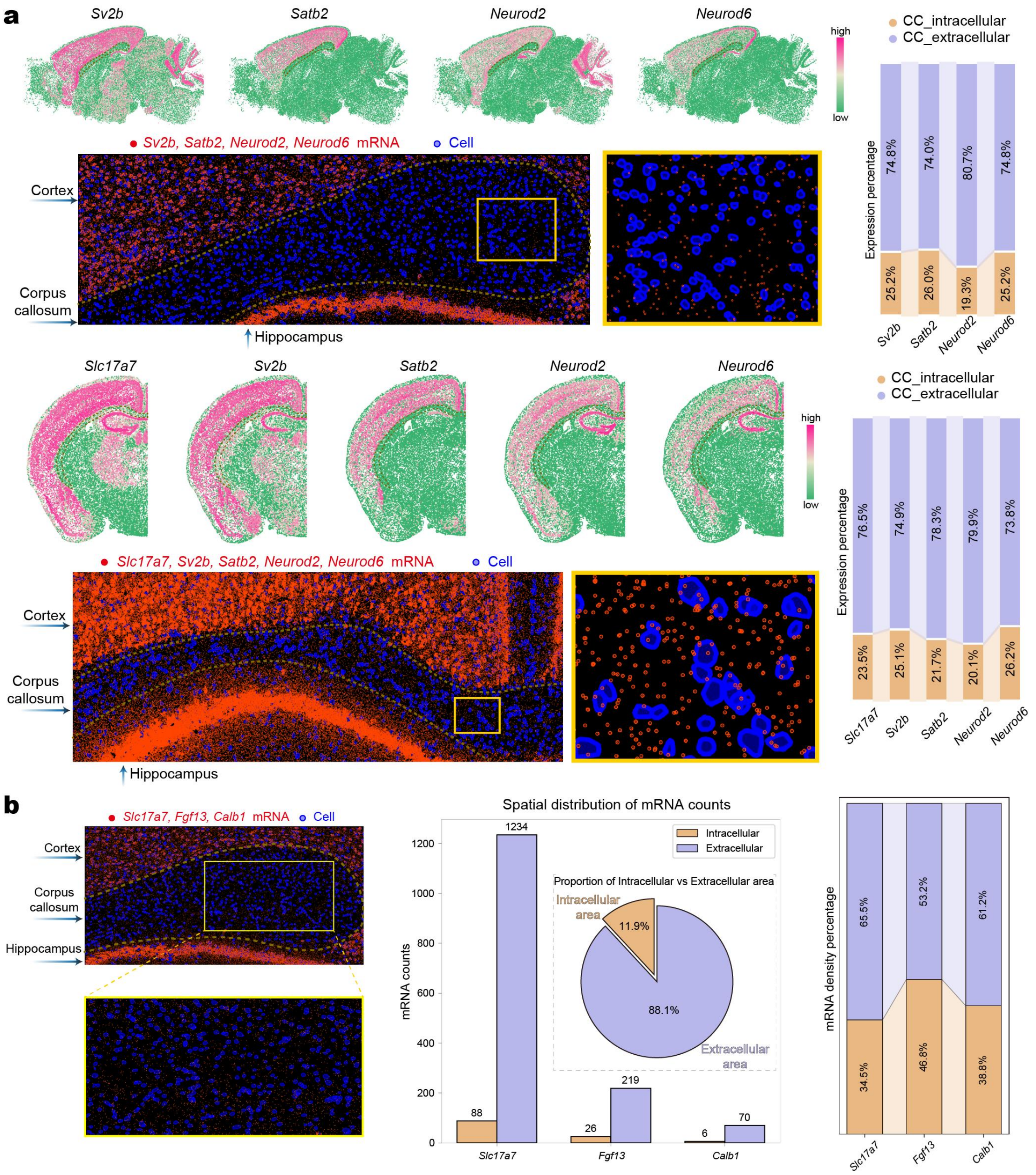


**Supplementary Figure S7: Comparison of intracellular and extracellular gene expression in the corpus callosum.**

**(a)** Differential gene expression between intracellular and extracellular compartments in the corpus callosum, as shown in sagittal (top) and coronal (bottom) MERFISH data. The examples include several marker genes primarily expressed in glutamatergic neurons, highlighting the localization of gene mRNA (red) and cells (blue) with enlarged views of the corpus callosum region. The stacked bar charts on the right illustrate the proportion of gene expression in intracellular (orange) and extracellular (purple) compartments.

**(b)** Distribution density of mRNA from cortical glutamatergic neurons in intracellular and extracellular compartments of corpus callosum. *Left*: Distribution of the corresponding mRNA (red) in the corpus callosum. *Middle*: Bar plot shows mRNA counts of three genes in intracellular (orange) and extracellular (purple) compartments of corpus callosum. Pie chart indicates the relative area proportion of intracellular (orange) and extracellular (purple) regions in the selected area from the left panel. *Right*: Stacked bar plot shows the distribution density of corresponding genes in intracellular (orange) and extracellular (purple) compartments of the corpus callosum.

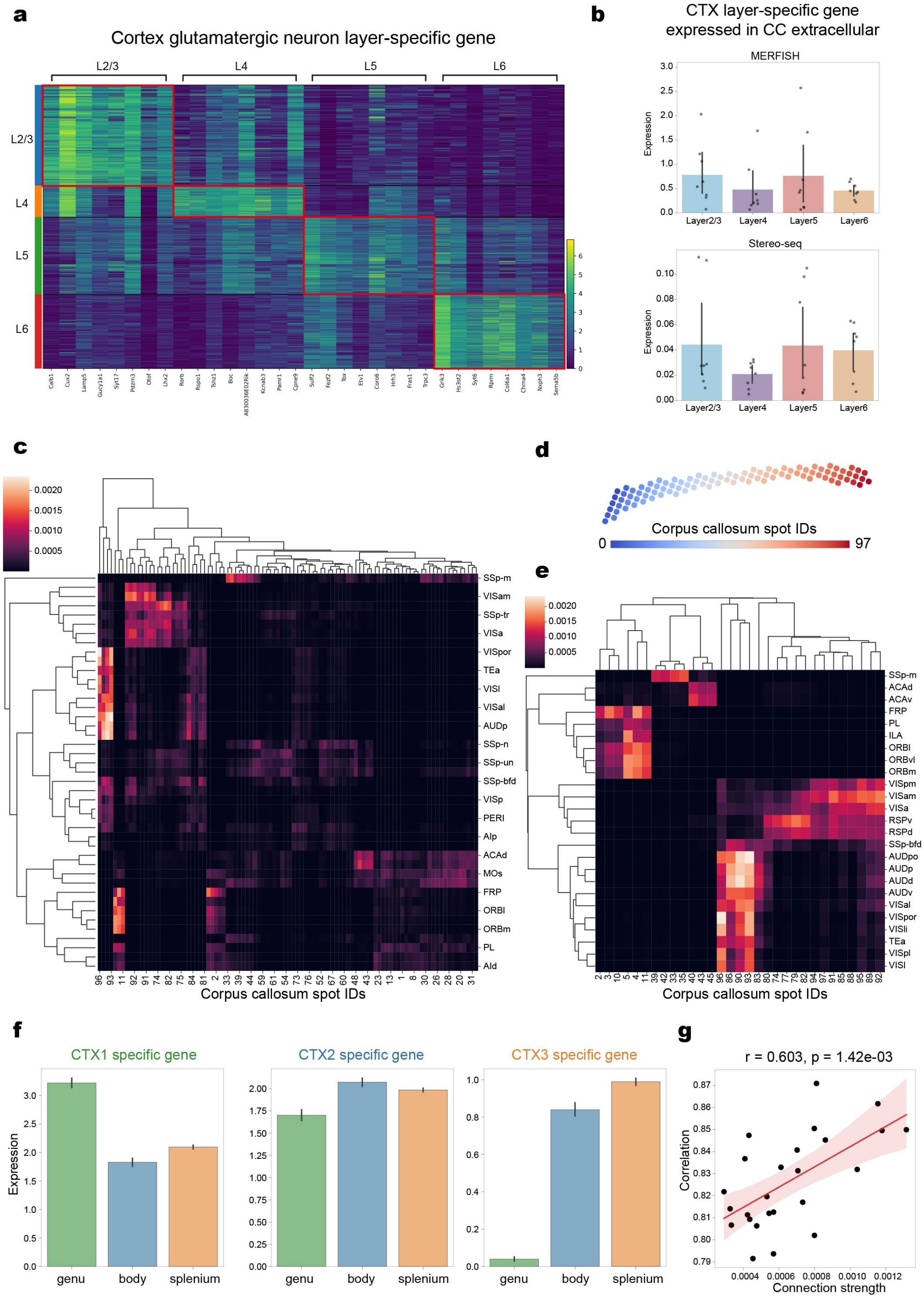


**Supplementary Figure S8: Relationship between cortical-corpus callosum connection strength and gene expression correlation.**

**(a)** The heatmap depicts the expression levels of genes enriched in specific cortical layers.

**(b)** Expression of cortical glutamatergic neurons layer-specific genes in intracellular and extracellular compartments within the corpus callosum. Bar plots present the average expression levels of cortical layer-specific genes in the intracellular and extracellular compartments of the corpus callosum, derived from MERFISH (top) and Stereo-seq (bottom) datasets. Error bars represent the 95% confidence intervals.

**(c)** The heatmap shows the connection strength from various cortical regions to each spot in the corpus callosum, ordered by hierarchical clustering. The x-axis represents the spot location numbers in the corpus callosum, as shown in (d), and the y-axis represents the cortical regions.

**(d)** Visualization of the spatial locations of corpus callosum spots (corpus callosum spot IDs), arranged sequentially along the corpus callosum.

**(e)** Similar to (c) but the weak connections are filtered (see **Methods**).

**(f)** Expression of cortical area-specific genes in extracellular compartments of the corpus callosum. The cortex and corpus callosum were divided into three compartments each according to cortico-callosal projections (**Fig. 2g**). Bar plots show expression levels of the top 5 differentially expressed genes (DEGs) from each cortical compartment within extracellular compartments of the corpus callosum. Error bars denote 95% confidence intervals. Similar to **Fig. 2h**, but based on the Stereo-seq data [3].

**(g)** Association between connection strength and gene expression correlation across cortical-callosal regions. Each data point corresponds to a cortical-callosal region pair (25 cortical regions×3 callosal regions). The x-axis shows connection strength between paired regions, while the y-axis shows their gene expression correlation. In the figure, $r$ represents the Pearson correlation coefficient of all points, and $p$ represents the corresponding p-value.

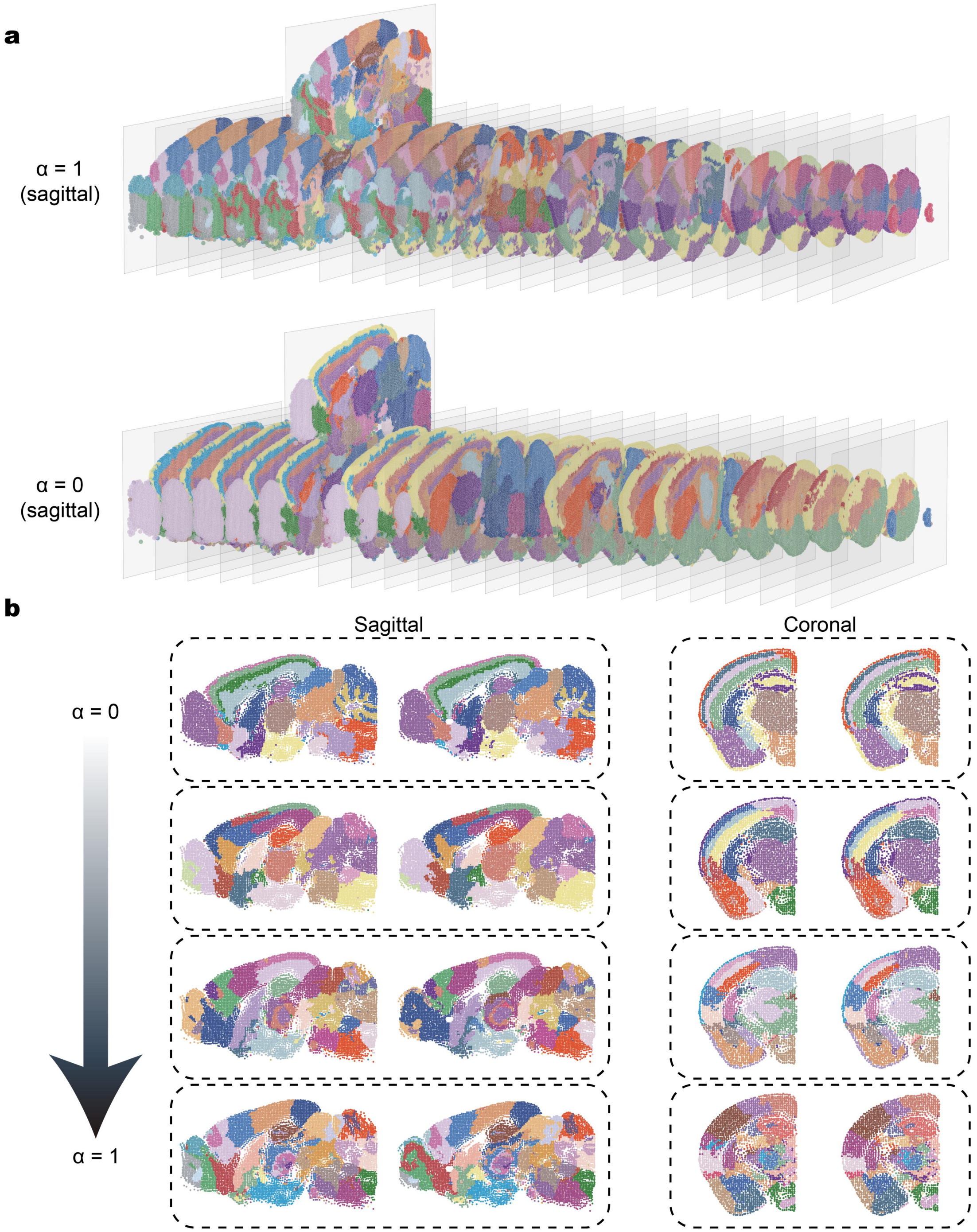


**Supplementary Figure S9: Clustering results of SpaCon under different α.**

**(a)** The 3D clustering results obtained by SpaCon under different α. Different colors represent distinct spatial domains identified by SpaCon.

**(b)** Clustering results of SpaCon under different values of the parameter α. Each row represents the clustering outcomes for a specific α value, with α increasing from top to bottom ($\alpha = 0$, $\alpha = 0.3$, $\alpha = 0.6$, $\alpha =$

1). The left panels show results from sagittal slices, and the right panels show results from coronal slices. Different colors represent distinct spatial domains identified by SpaCon.

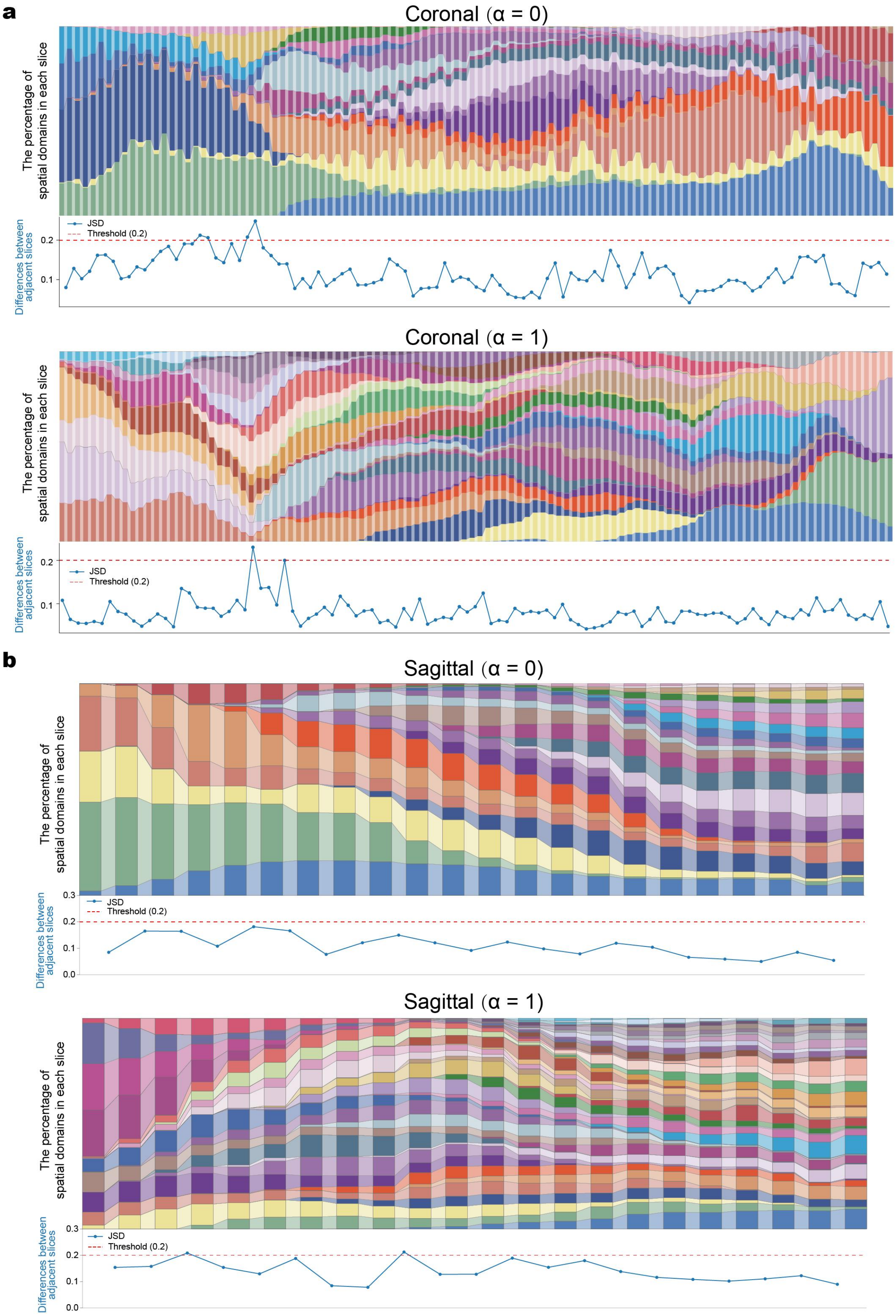


**Supplementary Figure S10: Evaluating batch effects of clustering results.**

**(a)** Stacked bar plots show proportions of spatial domains within each section, with color codes representing distinct domains. The stacked bar plots are arranged by original section order. Line plots depict Jensen-Shannon divergence (JSD) between adjacent sections, where the red dashed line (JSD=0.2) indicates the empirical threshold for negligible batch effects (JSD<0.2 suggests smaller inter-section variability). *Top*: Results with $\alpha=0$. *Bottom*: Results with $\alpha=1$.

**(b)** Similar to (a) but based on the sagittal MERFISH data.

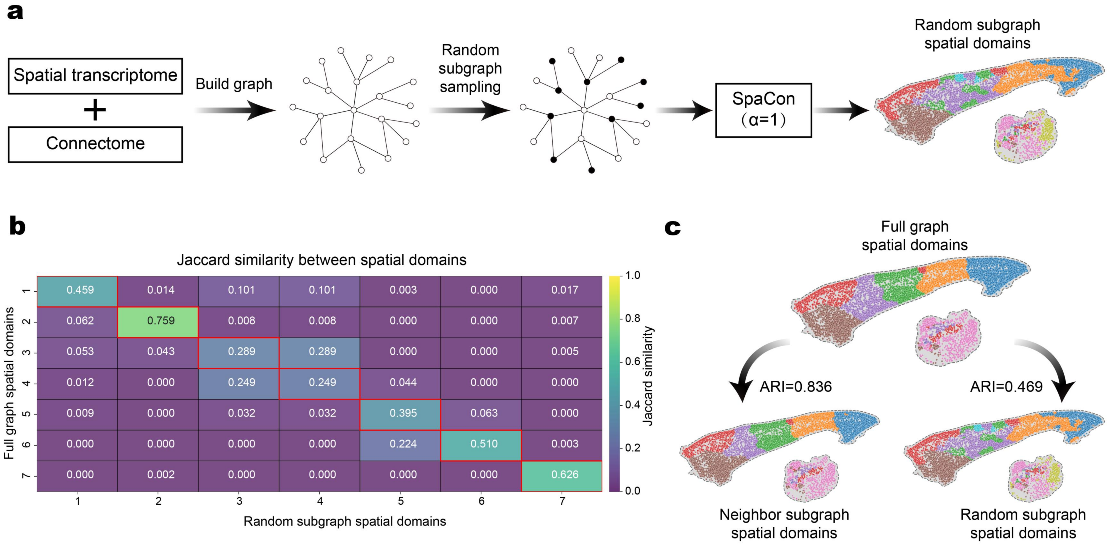


**Supplementary Figure S11: Results of different subgraph sampling methods.**

**(a)** Schematic diagram of random subgraph sampling. For the graph constructed by spatial transcriptomics and connectomics data, random sampling is first used to obtain subgraphs for batch model training and obtain the corresponding clustering results.

**(b)** Heatmap showing pairwise Jaccard similarity coefficients between spatial domains identified by full graph clustering (y-axis) and random subgraph clustering (x-axis). Color scale represents Jaccard similarity index values (intersection over union), with higher values indicating greater clustering consistency.

**(c)** The Adjusted Rand Index (ARI) between random subgraph spatial domains, neighbor subgraph spatial domains and full graph spatial domains. Here, ARI quantifies the similarity between clustering results.

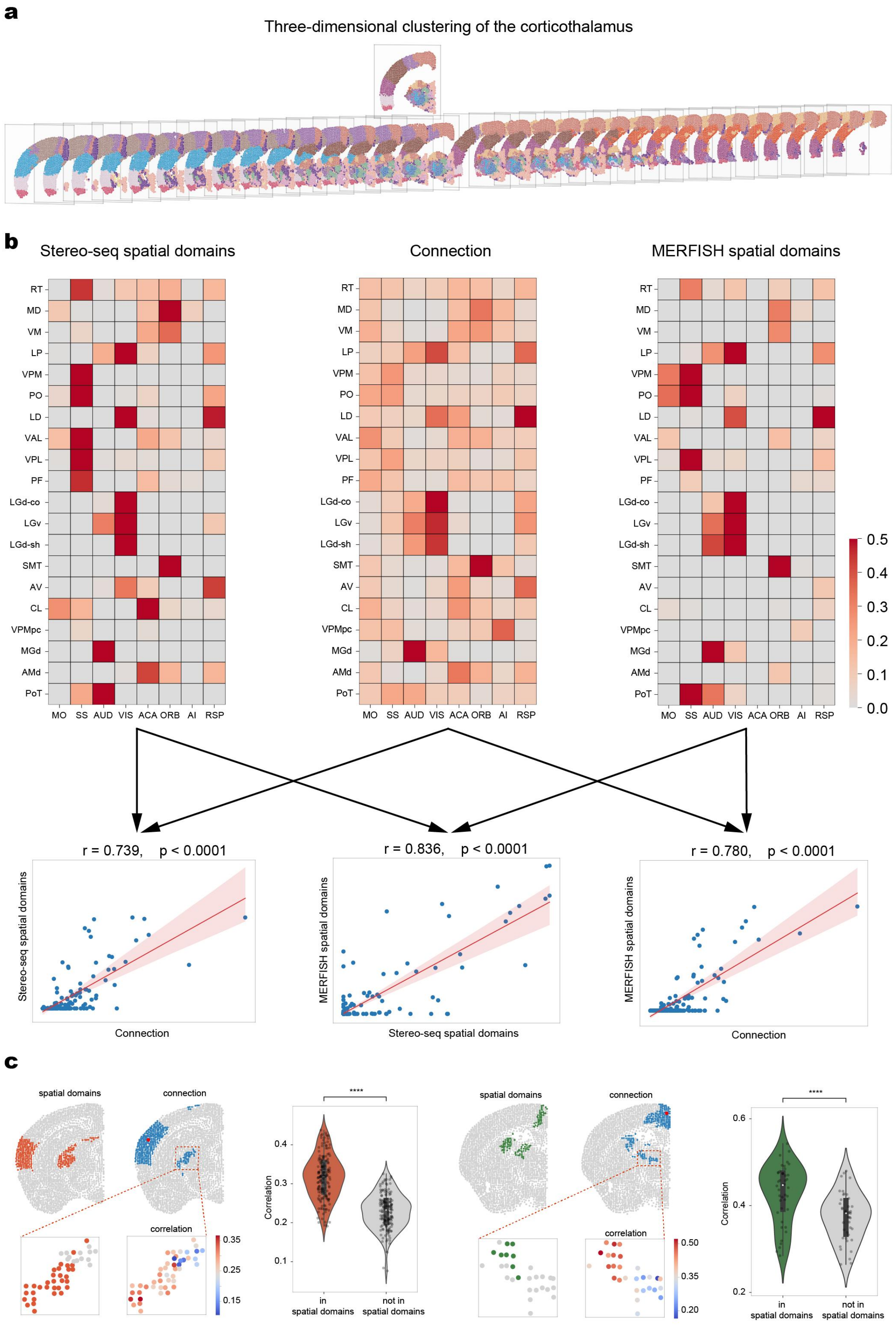
a
Three-dimensional clustering of the corticothalamus
b
Stereo-seq spatial domains
Connection
MERFISH spatial domains
RT
MD
VM
LP
VPM
PO
LD
VAL
VPL
PF
LGd-co
LGv
LGd-sh
SMT
AV
CL
VPMpc
MGd
AMd
PoT
MO SS AUD VIS ACA ORB AI RSP
0.5
0.4
0.3
0.2
0.1
0.0
r = 0.739, p < 0.0001
r = 0.836, p < 0.0001
r = 0.780, p < 0.0001
Stereo-seq spatial domains
Connection
MERFISH spatial domains
Stereo-seq spatial domains
MERFISH spatial domains
Connection
c
spatial domains
connection
correlation
0.35
0.25
0.15
****
Correlation
in spatial domains
not in spatial domains
0.50
0.35
0.20

**Supplementary Figure S12: Relationship between SpaCon clustering results and connectivity maps for corticothalamic regions.**

**(a)** Three-dimensional clustering results of the corticothalamic regions obtained by SpaCon (α=1). Different colors represent distinct spatial domains identified by SpaCon. All coronal slices are arranged along the rostro-caudal axis.

**(b)** The top three heatmaps display clustering results for corticothalamic regions from Stereo-seq [3] (*left*), corticothalamic connectivity-strength maps (*middle*), and MERFISH [2] clustering results (*right*). In the clustering heatmaps, color intensity represents the percentage of spots from each thalamic nucleus within each spatial domain associated with different cortical areas (see **Methods**). The connectivity heatmap represents the average connection strength between cortical and thalamic regions, based on the Allen Mouse Brain Connectivity Atlas [1]. The bottom three scatter plots illustrate the relationship between the clustering results and the connectivity maps. Each scatter plot corresponds to a pair of the top heatmaps. The red line represents the fitted regression line, with the shaded area indicating the 95% confidence interval.

**(c)** This figure presents supplementary results for two sets of spatial domains, similar to those shown in **Fig. 4b**. For each set, the coronal slice on the left shows the spatial domains identified by SpaCon, while the right panel displays the corresponding connectivity maps, where regions with connection strength greater than 0.001 to the highlighted red point are marked in blue. Magnified views illustrate the overlap between the connectivity maps and spatial domains. In the left magnified view, colored dots represent points within the spatial domain, whereas gray dots indicate points outside the domain. The right magnified view shows the Pearson correlation coefficient between the gene expression of each thalamic point and the red point in the cortex. The violin plots provide a statistical analysis of the gene expression correlations between the points within the spatial domain and those outside the domain (see **Methods**). ****: $p < 0.0001$.

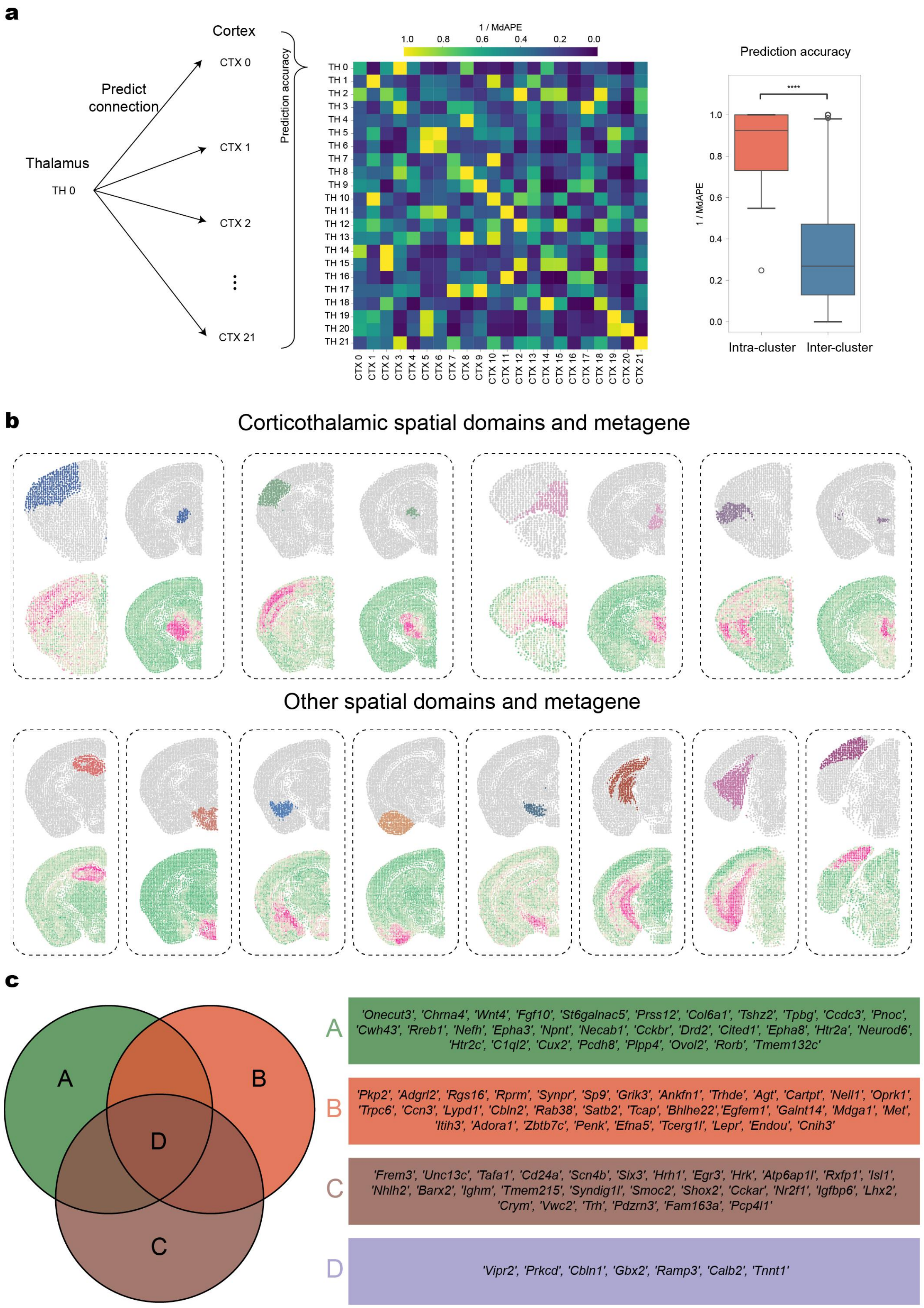


**Supplementary Figure S13: Additional results of metagenes of spatial domains identified by partial least squares (PLS) regression.**

**(a)** Predictive relationships of corticothalamic spatial domains based on SpaCon clustering. Similar to **Fig.4c-d**, but based on the coronal MERFISH data.

**(b)** The upper section shows spatial domains associated with corticothalamic connections (long-range connections) and the expression patterns of their metagenes. The lower section displays spatial domains without long-range connections and corresponding metagenes. In each section, the first row presents the spatial domains identified by SpaCon. The second row shows the expression levels of a set of metagenes associated with these spatial domains, as identified by PLS regression.

**(c)** Genes associated with the spatial domains identified in **Fig. 4f**. The Venn diagram on the left shows the overlap of genes identified by PLS across three spatial domains, with A, B, and C representing genes specific to each domain, and D representing genes common to all three domains. The gene names corresponding to each category are listed on the right.

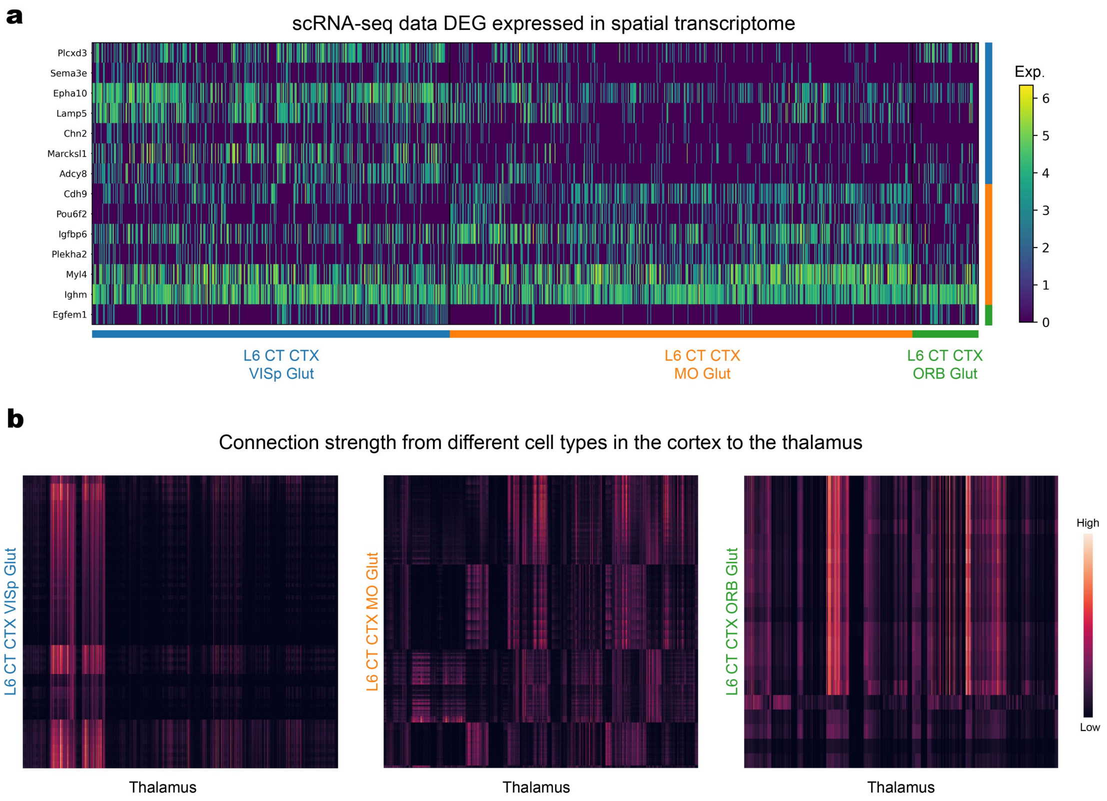


**Supplementary Figure S14: Additional results of SpaCon for cell subtype classification.**

**(a)** The heatmap shows the expression of these genes in each cell type of the spatial transcriptome. Among them, these genes are DEGs of each cell type obtained based on scRNA-seq data (**Fig. 6e**). L6 CT CTX VISp Glut: Corticothalamic glutamatergic neuron subtype located in layer 6 of cortical VISp. L6 CT CTX MO Glut: Corticothalamic glutamatergic neuron subtype located in layer 6 of cortical MO. L6 CT CTX ORB Glut: Corticothalamic glutamatergic neuron subtype located in layer 6 of cortical ORB.

**(b)** The heatmap quantifies connection strength from three defined cellular subtypes to thalamic (**Fig. 6f**). The thalamic cells in the three heatmaps are arranged in the same order. The differences between the three heatmaps indicate that they project to different thalamic regions.

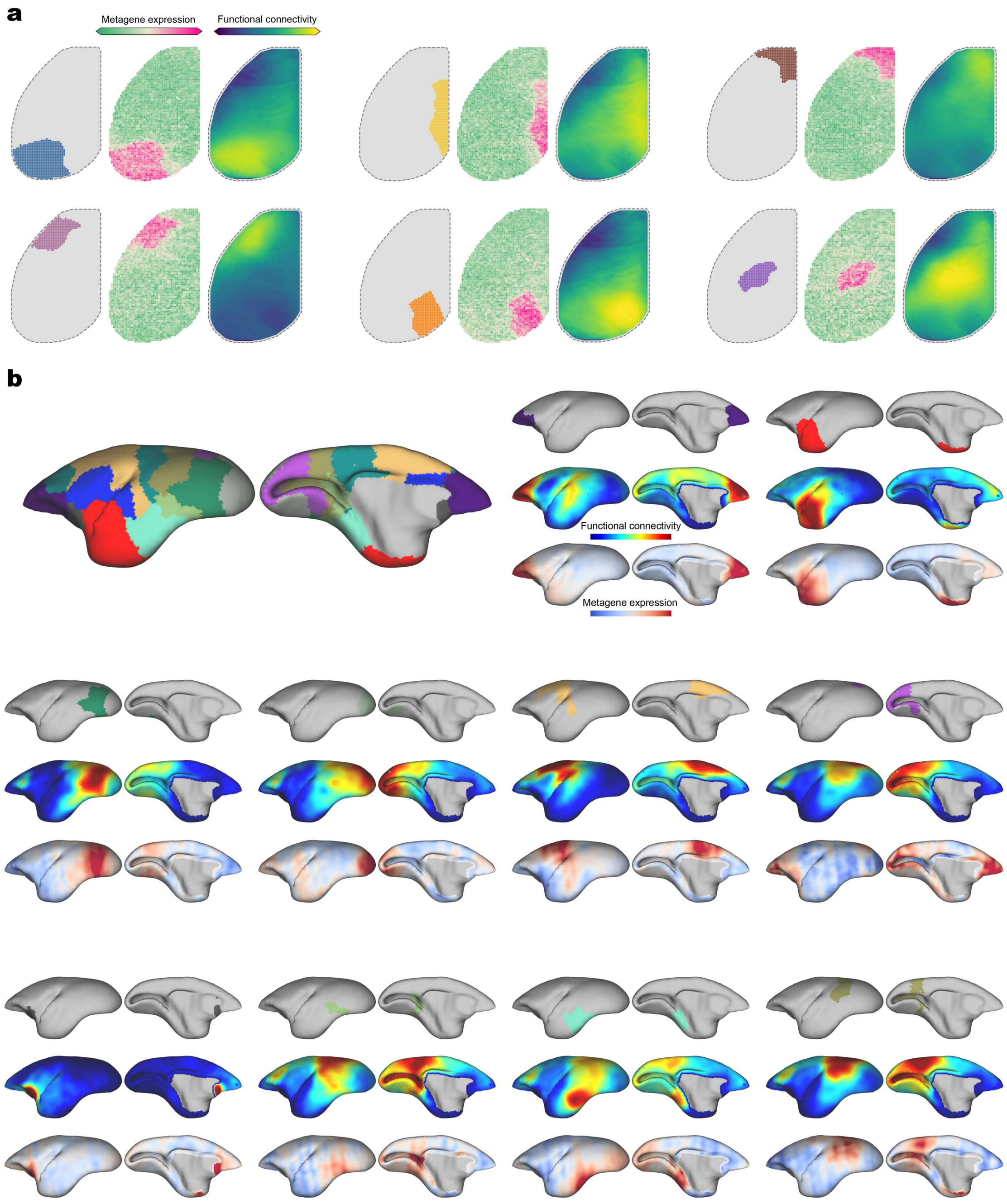


**Supplementary Figure S15: Additional results of SpaCon application to wide-field calcium imaging and fMRI data.**

**(a)** Results of SpaCon applied to wide-field calcium imaging and spatial transcriptomics (**Fig. 6a**). Each set of results shows the SpaCon spatial domain, functional connection strength of the spots from each domain across the entire cortical region, and cortical expression of a set of metagenes identified by PLS within each spatial domain.

**(b)** Results of SpaCon applied to fMRI and spatial transcriptomics (**Fig. 6d**). Each set of results shows the SpaCon spatial domain, functional connection strength of the spots from each domain across the entire cortical region, and cortical expression of a set of metagenes identified by PLS within each spatial domain.

### Supplementary Figures References
